# Cell-to-cell variability in AMPK activation reveals autonomous cycles in cellular energy balance

**DOI:** 10.1101/780023

**Authors:** Nont Kosaisawe, Breanne Sparta, Michael Pargett, Carolyn Teragawa, John G. Albeck

## Abstract

Cellular metabolism can be reconfigured to balance nutrient processing (catabolism) and cellular demands (anabolism). However, the kinetics of reconfiguration within individual mammalian cells, and heterogeneity between cells, have remained unexplored. Using live-cell imaging, we investigate the kinetics of bioenergetic adaptation in individual cells. In response to acute inhibition of oxidative phosphorylation, AMPK substrates are phosphorylated bimodally, identifying cells in different underlying states of anabolism and catabolism. Manipulation of glycolysis, insulin/Akt signaling, or protein synthesis shifts the distribution of these states. Long-term lineage analysis confirms that this cellular energy balance cycles over time within individual cells, independently of the cell division cycle. We further demonstrate that AMPK inhibits the ERK and mTOR cell growth signaling pathways specifically when anabolism is in excess. Our results reveal dynamic fluctuations of energetic balance, establish distinct time scales of cellular energetic control, and open opportunities for more precise prediction and control of cellular metabolic functions.

## Introduction

An essential feature of cellular metabolic regulation is the capability to adjust metabolite flux to support functions including cell growth and division. This adaptability, sometimes termed metabolic plasticity, has been observed between cell types and tissues (Konagaya et al., 2017; Simões et al., 2015) and contributes to the development and metastasis of cancer (DeBerardinis et al., 2008; Vander Heiden and DeBerardinis, 2017). However, it remains uncertain to what extent individual cells vary over time in their metabolic configuration, or differ from cell to cell. In part, this uncertainty arises from the use of bulk analysis methods with low temporal resolution, while it is clear that both the cell cycle and various signaling processes proceed asynchronously within any population of cells (Wollman, 2018). Temporal fluctuations in metabolic function are best understood in the model organism *S. cerevisiae*, in which metabolic cycles proceed in synchrony with the cell cycle under certain conditions (Cai and Tu, 2012). Such cycles may also occur independently of the cell cycle (Silverman et al., 2010), or regulate passage through cell cycle checkpoints (Papagiannakis et al., 2017). In mammalian cells, metabolic cycling has been observed in concert with the cell cycle (Ahn et al., 2017) or with circadian rhythms (Bass and Takahashi, 2010). However, the question of whether individual mammalian cells possess independent metabolic cycles remains unresolved.

A key control point for metabolic plasticity is the balance between the use of glycolysis and oxidative phosphorylation (OxPhos) for production of ATP. A prominent example of such regulation is the Warburg effect, also known as aerobic glycolysis, in which the rate of glycolysis is enhanced even in the presence of adequate oxygen, a shift that enables tumor cells, and other rapidly proliferating cells such as activated lymphocytes, to efficiently produce biosynthetic intermediates needed for cell proliferation (Gatenby and Gillies, 2004)(Vander Heiden et al., 2009). Importantly however, the choice between glycolysis and OxPhos is not exclusive or discrete, and these processes may be controlled independently (Vander Heiden and DeBerardinis, 2017). Because of the centrality of this balance, sophisticated assays have been developed to measure the capacity of these two pathways (Fan et al., 2013; Mookerjee et al., 2017a). However, these assays require thousands of cells and cannot report on individual cell variation in metabolic balance.

A major challenge in assessing cellular metabolic state is the fact that reaction fluxes are not directly observable by simple measurement of any one metabolite. Additional information is required, which can be obtained by tracing the fates of labeled metabolites (Fendt et al., 2013), or by fitting kinetic models to parallel measurements of enzyme concentrations (Hackett et al., 2016). Both of these strategies require fairly comprehensive measurements to ensure that all important routes for flux are accounted for. Such analyses require a large samples and are currently infeasible in single cells. Another approach is to monitor the response to defined perturbations, tracking the immediate change in key metabolites to gauge the impact on the metabolic network. For example, electron transport chain (ETC) inhibitors, such as oligomycin, have been applied to measure the production of ATP by oxidative phosphorylation, by tracking the acute change in oxygen consumption (Buttgereit and Brand, 1995). This approach requires frequent sampling and an observable that can be monitored in live cells as they respond to the perturbation. Such observables can either be metabolites such as lactate or oxygen, or endogenous proteins that serve as cellular sensors for metabolic status. One such sensor is AMP-activated protein kinase (AMPK), a regulatory hub that coordinates metabolism and cell growth. AMPK is a heterotrimeric protein that directly binds ATP, ADP, and AMP, and whose kinase activity is increased upon binding by ADP and AMP. In response to decreased cellular energy charge (AMP and ADP relative to ATP)(Hardie and Hawley, 2001; Oakhill et al., 2011), it phosphorylates an array of substrates to enhance catabolism (e.g., glycolysis, fatty acid oxidation, and autophagy) and suppress anabolism (e.g., synthesis of proteins and DNA) (Hardie, 2014). Elegant biochemical and structural studies have elucidated the basis for this sensing by the trimeric AMPK complex at the protein level (Gowans et al., 2013; Xiao et al., 2011). At the cellular level, AMPK activity and phosphorylation, or phosphorylation of its effectors such as acetyl-CoA carboxylase (ACC), are frequently used as indicators of low energy charge.

Recently, using a live-cell AMPK activity reporter (AMPKAR2), we discovered a high degree of variability in AMPK activity in cultured epithelial cells (Hung et al., 2017a). Treatment with ETC inhibitors (ETCi) induced cycles of AMPK activation with a period of 3-5 hours, indicating that, at least under certain conditions, the metabolic activity of individual cells can cycle in a rhythm independent from the cell cycle. We also detected variability from cell to cell in the initial induction of AMPK activity, suggesting that there may be pre-existing metabolic states that predispose ETCi-resistant and ETCi-sensitive responses. Potentially, these measurements of initial response between cells could parallel, at a single cell level, a widely used assay for the the rate of ATP production by OxPhos, in which the change in oxygen consumption is measured during ETCi treatment (Buttgereit and Brand, 1995; Mookerjee et al., 2017a). In principle, a strong AMPK response to ETCi, would indicate a dependence on OxPhos for ATP production. Conversely, cells with adequate capacity to generate ATP by glycolysis would not experience a loss in energy charge and activation of AMPK upon ETCi treatment. Therefore, this experimental strategy may provide a tool to distinguish differences in metabolic configuration with single-cell resolution. However, such an approach requires validation of the underlying changes in ATP metabolism, as AMPK can respond to a variety of changes in metabolic status (Lin and Hardie, 2017; Zhang et al., 2017). To address this ambiguity, reporters for intracellular ATP concentration (Ateam1.03) (Imamura et al., 2009b), or ADP/ATP ratio (PercevalHR)(Tantama et al., 2013a) provide useful tools to make independent measurements of cellular ATP metabolism (Figure 1A).

**Figure 1.**
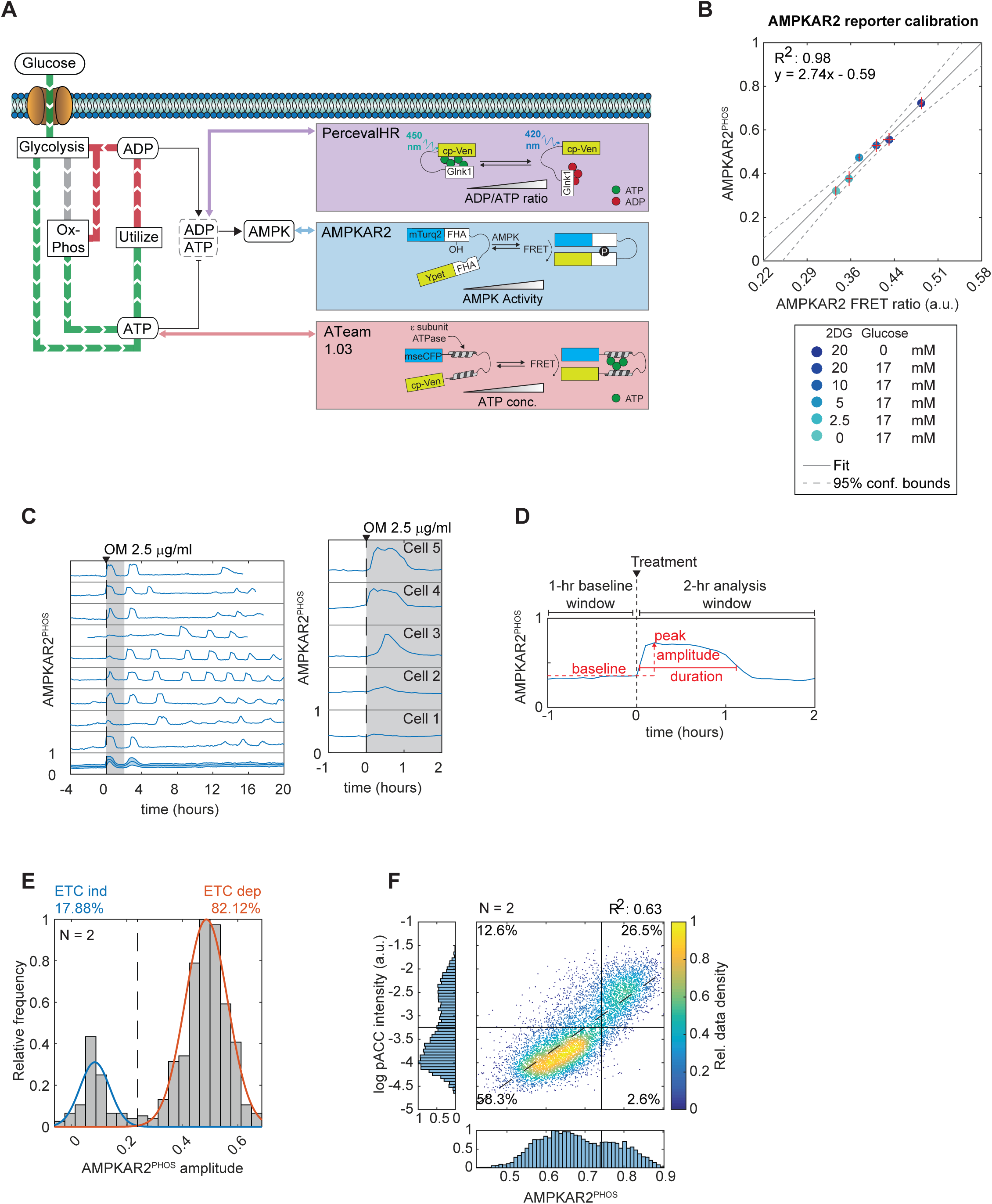
Quantitative AMPK reporter measurements identify subpopulations of cells resistant to ETC inhibition stress. A: Schematic of ATP metabolism and the reporters used in this study. AMPKAR2 indicates AMPK kinase activity, PercevalHR reports intracellular ADP/ATP ratio, and ATeam1.03 reports intracellular ATP concentration. All these sensors operate at minute-scale resolution, providing an immediate readout of metabolic changes after perturbation. B: Scatter plot of the correlation between FRET ratio of AMPKAR2 reporter as measured by live-cell microscopy and its phosphorylation status as measured by phos-tag gel electrophoresis under the same conditions. A range of different AMPK activities were induced by varying glucose and 2-deoxyglucose (2-DG). Error bars represent standard errors of the mean (SEM) from at least two different experiments. The solid line represents a fitted linear model, and the dashed lines show 95% confidence bound. This fitted equation is used throughout the study to report all AMPKAR2 FRET measurements as the fraction of AMPKAR2 phosphorylated [x – AMPKAR2 FRET ratio; y - AMPKAR2^PHOS^]. C: Left panel: representative AMPKAR2^PHOS^ measurements for cells grown in 17 mM glucose, after treatment with 2.5 µg/ml oligomycin. Each subplot represents a single cell measurement, with population average and interquartile range in the bottom subplot. The shaded area shows the 2-hour time window of the data used for analysis. Right panel: enlargement to show details of acute AMPKAR2^PHOS^ responses. D: Schematic of AMPKAR2 pulse parameterization. Peak activity was defined as the local maxima value within 2 hours after perturbation; baseline was defined as the average of AMPKAR2 activity for one hour before treatment. Amplitude was calculated by subtraction of baseline from peak. Pulse duration is defined as full width at half-maximum. E: Histogram of AMPKAR2^PHOS^ amplitudes after treatment with 2.5 µg/ml oligomycin. Blue and orange lines are fitted Gaussian distributions. The dashed line is defined by the intersection between distributions and used as the cutoff for ETC-ind and ETC-dep cells. F: Scatter plot of single-cell measurements of AMPKAR2^PHOS^ and phospho-ACC staining intensity in MCF10A cells treated with 2.5 µg/ml oligomycin. AMPKAR2^PHOS^ was measured by live-cell microscopy, and phospho-ACC values were calculated for the same cells after fixation and staining (see Methods for details). The dashed line shows a fitted linear function. Numbers indicate the percentage of cells in each quadrant.

In this study, we investigated how cell-to-cell differences in AMPK responses to ETCi report on individual cell metabolic behavior, using multiple genetically encoded reporters and supporting assays to track acute adaptive metabolic responses. We demonstrate that epithelial cell lines contain isogenic cells with both high and low glycolytic capacity relative to ATP consumption, whose frequencies are modulated by insulin-stimulated glucose uptake and protein translation rate. Transition from one configuration to the other occurs on the timescale of hours to days, but is not closely linked to the cell cycle. These findings establish a new layer of plasticity in mammalian cellular energetic configuration at the level of individual cells.

## Results

### Quantitative AMPK reporter measurements identify subpopulations of cells resistant to ETCi stress

AMPK responds to reduced energy charge, and cell-to-cell variation in AMPK activity in response to ETCi may thus reflect each cell’s reliance on OxPhos for ATP production. However, this variation could also arise from other sources, including non-linearity or non-specificity in the reporter readout. We investigated these alternatives using the non-tumor epithelial cell line MCF10A, a system that responds with metabolic changes to physiological signals including insulin, growth factors, and attachment (Grassian et al., 2011; Hung et al., 2017a; Schafer et al., 2009). To quantitatively relate AMPKAR2 reporter signals to cellular AMPK activity, we calibrated the FRET ratio measured by microscopy to AMPKAR2 phosphorylation status using Phos-tag electrophoresis (Figure S1A-B). AMPKAR2 FRET ratio was linearly correlated with AMPKAR2 phosphorylation fraction across the full range of metabolic conditions tested (Figure 1B). Under conditions of extreme metabolic stress (glucose starvation or treatment with the glycolytic inhibitor 2-deoxy glucose) only ∼75% of AMPKAR2 sensor was phosphorylated, indicating that the reporter was not saturated. In cells cultured in full growth medium, ∼30% of AMPKAR2 was phosphorylated, reflecting detectable AMPK activity even under high nutrient conditions. Because phosphorylation of AMPKAR2 parallels the phosphorylation of endogenous AMPK substrates (Tsou et al., 2011), we report all subsequent AMPKAR measurements as the calibrated fraction of AMPKAR2 phosphorylated, referred to as “AMPKAR2^PHOS^”.

We next quantified the heterogeneity in AMPK substrate phosphorylation immediately after ETCi treatment. The magnitude of AMPK activity for each cell was quantified as the baseline-to-peak amplitude of AMPKAR2^PHOS^ within 2 hours following oligomycin treatment (Figure 1C-D). In cells cultured in high glucose, this metric was distributed bimodally, with ETC-dependent (ETC-dep) cells distributed around a mode of ∼0.5 and ETC-independent (ETC-ind) cells at a mode approximately 6-fold lower (Figure 1E). The ETC inhibitors antimycin and rotenone produced similar distributions (Figure S1D), indicating that these responses are a property of the regulatory network rather than the specific form of inhibition. ETC-ind cells were not a result of sub-saturating inhibitor, as a maximal effect was achieved at 0.3 µg/ml oligomycin, and was not further increased by a 30-fold higher concentration (Figure S1E). Measurements of oxygen consumption rate confirmed that oligomycin at 1 µg/ml yields a suppression equivalent to the maximal achievable effect in the presence of all three ETCi (Rotenone + Antimycin A + Oligomycin) (Figure S1F). Inference of ATP production (Mookerjee et al., 2017a) confirmed a nearly complete switch from OxPhos to glycolysis during ETCi treatment (Figure S1G), although glucose uptake was increased only weakly by ETCi (Figure S1H), suggesting a redistribution of glycolytic metabolite flux. Metabolomic analysis confirmed that oligomycin treatment induced a profound suppression of the TCA cycle intermediates citrate, aconitate, isocitrate, and α-ketoglutarate, and a more heterogeneous shift in glycolytic intermediates (Figure S2). We conclude that the lack of AMPK activity in ETC-ind cells is not an artifact of incomplete inhibition, and that ETC-ind cells represent a subpopulation with a metabolic configuration inherently distinct from ETC-dep cells.

To corroborate that heterogeneity in AMPKAR2^PHOS^ reflects differences in endogenous AMPK substrate phosphorylation, we co-stained cells with an antibody specific for ACC phosphorylated at Ser-79 (pACC), a canonical AMPK substrate, immediately after recording their live-cell AMPKAR2^PHOS^ amplitude by live-cell imaging (Figures 1F and S1C). pACC staining was bimodal and correlated with AMPKAR2^PHOS^ (R^2^=0.63), with >80% of cells either double-positive or double-negative. Thus, both reporter and endogenous substrate reveal the existence of ETCi-dep and ETCi-ind cells within the population.

### Single-cell AMPK responses to ETCi report changes in energy charge

Having established concordance of AMPKAR2 with endogenous AMPK substrates, we next asked whether variation in AMPK activity results from heterogeneity in cellular energy charge (Hardie, 2011; Hardie et al., 2012), or from other factors modulating AMPK. To track energy charge, we performed a similar analysis with the ADP/ATP reporter Perceval HR (Berg et al., 2009; Tantama et al., 2013a), which reports intracellular ADP/ATP ratio as a spectral shift in mVenus excitation, a ratio we refer to as Perceval^EX^. Similar to AMPKAR activity, we also observed heterogeneity in the immediate response of Perceval^EX^ following ETCi (Figure 2A). However, unlike AMPK activity, Perceval^EX^ lacked two distinct modes. Under continuous exposure to oligomycin, we observed pulses of Perceval^EX^ 1-2 hours in duration, interspaced by 2-4 hours, similar to AMPKAR2^PHOS^ in timing but more variable in amplitude. Immunofluorescence staining of pACC was strongly correlated with Perceval^EX^, with agreement of pACC staining and Perceval^EX^ responses in ∼80% of cells (Figures 2B and S3A). However, the distinction between high- and low-Perceval^EX^ cells was not sharp, and cells at intermediate Perceval^EX^ values were distributed between high- and low-pACC subpopulations, consistent with findings that factors in addition to energy charge can influence AMPK activity (Hardie et al., 2016; Hawley et al., 2005; Woods et al., 2005; Zhang et al., 2017). Nonetheless, the overall strong correlation between pACC and Perceval^EX^ indicates that energy charge is the primary factor shaping AMPK activity under these conditions, and that variation in energy charge can account for the observed cell-to-cell variation in AMPK activation.

**Figure 2.**
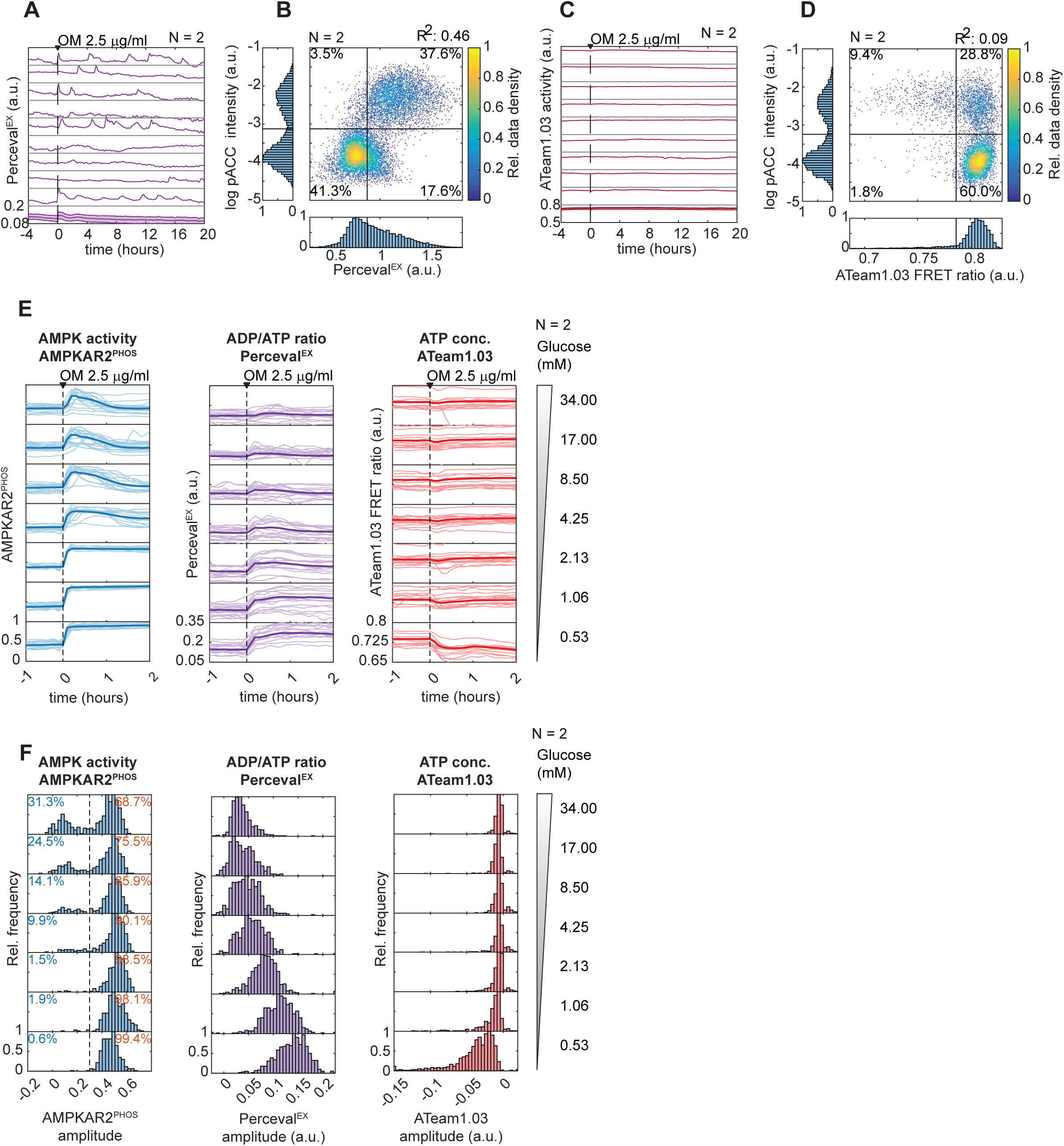
Divergent AMPK responses to ETCi indicate changes in energy charge despite stable ATP concentration. A and C: Representative single-cell activities of PercevalHR (A) or ATeam1.03 (C) after treatment with 2.5 µg/ml oligomycin. Each subplot represents a single cell measurement, with the population average and interquartile range shown in the bottom subplot. B and D: Scatter plots of single-cell measurements of PercevalHR activity (B) or ATeam1.03 activity (D) with phospho-ACC staining intensity in MCF10A cells treated with oligomycin 2.5 µg/ml. Numbers indicate the percentage of cells in each quadrant. R^2^ values are shown for linear fits to the data. E: Sample single-cell activities of each reporter after cells were cultured in media containing glucose as indicated and then treated with 2.5 µg/ml oligomycin. F: Histograms of response amplitude of each reporter, calculated as in Figure 1.

We next used the FRET-based ATP sensor ATeam 1.03 (Imamura et al., 2009) to track intracellular ATP concentrations under the same conditions. Strikingly, following oligomycin treatment, we were unable to detect any change in ATP level (Figure 2C), nor any pulsatile characteristics as observed for AMPKAR2^PHOS^ or Perceval^EX^. To confirm that the lack of ATeam response is not a result of out-of-range ATP concentration, we treated these cells with oligomycin in the absence of glucose, which resulted in an immediate and sharp decline in ATeam signal (Figure S3C), followed within 4 hours by visible cell death. We confirmed this result using bulk ATP assays, which detected no ETCi-induced change in ATP at 17 mM glucose, but a >90% decrease upon ETCi treatment in the absence of glucose (Figure S3D). When ATeam cells were co-stained with pACC, we observed that the rare low-ATeam cells (∼10%) were predominantly pACC-positive, as would be expected for cells with low ATP levels (Figures 2D and S3B). These results suggest that ATeam accurately reports ATP levels within MCF10A, and that cytoplasmic ATP abundance remains stable during OxPhos inhibition. Altogether, our reporters indicate that ATP, ADP/ATP ratio, and AMPK activity are interrelated, as expected, but not strictly proportional within individual cells.

To further characterize the relationship between energy charge and AMPK responses, we quantified AMPKAR2^PHOS^, Perceval^EX^, or ATeam responses following oligomycin treatment under varying concentrations of glucose (Figure 2E-F). AMPKAR2^PHOS^ responses remained bimodal across all conditions, with the frequency of ETC-ind cells decreasing from >20% of cells at 17 mM glucose (standard MCF10A culture conditions) to 8-9% at 4.25 mM glucose (an intermediate physiological concentration) and falling to <1% at lower glucose concentrations. In contrast, Perceval^EX^ was distributed unimodally in each condition, with a mean that increased gradually as glucose concentration was lowered. ATeam responses were detectable only at concentrations of glucose <1 mM. Together, these data are consistent with a model in which the absolute concentration of ATP is maintained at a nearly constant level, provided that glycolysis can operate at a sufficiently high rate. Rapid equilibration of ATP with ADP and AMP prevents a large drop in absolute ATP levels but allows a significant shift in ADP/ATP and AMP/ATP ratios, which are detected sensitively by AMPK (Hardie et al., 2012). Our results indicate that these ratios vary substantially from cell to cell, and are sufficient to induce AMPK activity in some cells (ETC-dep), but not others (ETC-ind). The bimodality observed in AMPK activity, but not ADP/ATP ratio, implies that phosphorylation of AMPK substrates occurs in an ultrasensitive manner when energy charge and AMPK activation cross a threshold, which may also vary from cell to cell.

### AMPK responses to ETC inhibition reveal cycling long-term metabolic configurations

The subpopulations defined by ADP/ATP and AMPK responses to ETCi could be relatively static, or could change as a function of other cellular processes. We investigated the cell cycle as one potential source of variability, recording single-cell profiles of AMPKAR2^PHOS^ over a 27-hour period. More than 11,000 cell profiles were aligned by the time of mitosis, allowing cell age to be used as a proxy for cell cycle phase (Figure 3A)(Sigal et al., 2006a). Both ETC-ind and ETC-dep AMPK responses were evenly distributed throughout the cell cycle. Similarly, young and old cells were randomly distributed when profiles were ordered by the strength of AMPKAR2^PHOS^ response to oligomycin (Figure 3B). The only systematic relationship between AMPKAR2^PHOS^ and the timing of mitosis was a decrease in the average amplitude of response for cells that had divided >20 hours earlier (Figure 3C). As the average cell cycle length in this population was 22 hours, this trend may reflect a lower average response in cells that are quiescent or cell cycle-arrested. Overall, our data rule out cell cycle phase as a major determinant of variable AMPK responses. We also failed to detect any strong correlation between basal AMPKAR2^PHOS^ and AMPKAR2^PHOS^ amplitude (Figure 3D).

**Figure 3.**
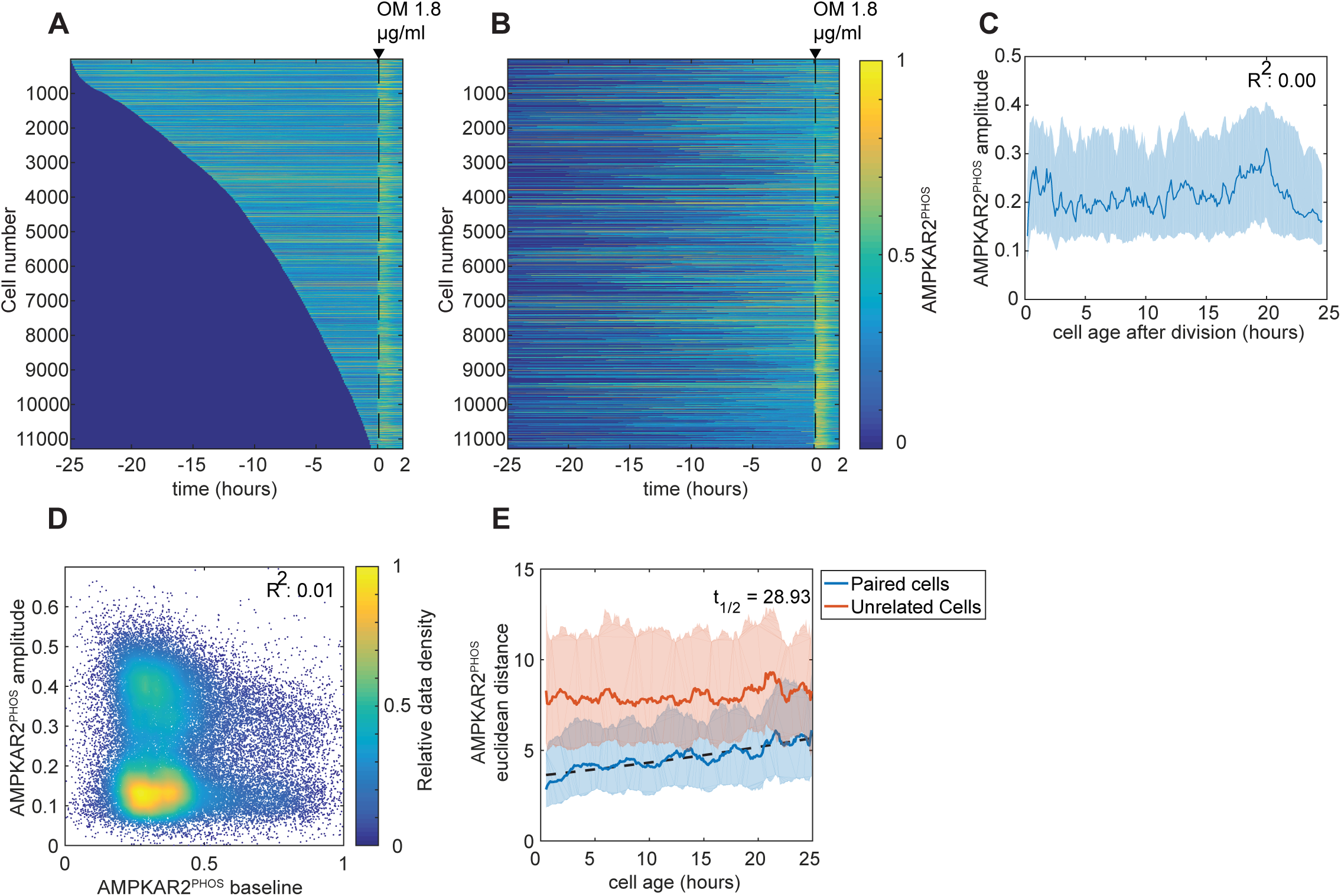
AMPK responses to ETC inhibition reveal fluctuating metabolic configurations. A and B: Heatmaps of AMPKAR2 activity in individual cells. MCF10A-AMPKAR2 cells were imaged for 24 hours before treatment with 1.8 µg/ml oligomycin. Each horizontal line represents a single cell’s AMPKAR2^PHOS^ profile (represented by color), beginning with its most recent cell division and ending 2hr after oligomycin treatment. Cells were sorted either by the time of their last division (A) or by the amplitude of AMPKAR2 response upon oligomycin treatment (B). Analysis contains >11,000 individual cells. C: Line plot showing the relationship between a cell’s age at the time of oligomycin treatment and its recorded AMPKAR2^PHOS^ amplitude response. Values were calculated for bins of 6 minutes. D: Scatter plot of baseline AMPKAR2^PHOS^ (prior to oligomycin treatment) and corresponding AMPKAR2^PHOS^ amplitude for each cell following oligomycin (1.8 µg/ml). Colormap represents relative data density. E: Comparison of AMPKAR2^PHOS^ responses in sister cells. Dissimilarity of the oligomycin responses between the sisters of each division was calculated using time-series Euclidean distance, and plotted as a function of the time between the last division and oligomycin treatment. Dissimilarity between pairs of unrelated cells of the same age is shown as a control (red line). Solid lines represent effect size, and the shaded areas represent interquartile percentile after bootstrapping. Dashed line represents a fitted exponential function for the decay of sister cell similarity over time. See STAR Methods for details.

Using the same dataset, we also performed a sister-cell analysis, in which we compared the ETCi responses of both daughter cells of each cell division as a function of their age (time between division and treatment with oligomycin). Sister cell pairs with ages less than 2 hours showed a difference significantly lower than random pairs of cells, indicating that the tendency toward ETC-ind or ETC-dep response is a heritable property that can persist for at least several hours (Figure 3E). The similarity in AMPKAR2^PHOS^ between daughters decayed gradually, such that it approached the level of unrelated cells with a half-life of ∼29 hours. Such patterns of decaying correlation have been observed in other processes, including cell death and signaling pathways (Spencer et al., 2009; Strasen et al., 2018), where they reflect shifts in protein or gene expression complement of each cell that alter cellular phenotype on the scale of 1-2 days (Sigal et al., 2006b). The calculated half-life value therefore provides an upper bound on the time needed for an individual cell to shift from ETC-dep to ETC-in, or vice versa, implying that cycling between these two states occurs regularly within the cell population.

### The balance between glycolysis and protein translation regulates AMPK heterogeneity

We next investigated factors involved in switching between ETC-ind and ETC-dep states. In principle, the difference between ETC-ind and ETC-dep cells lies in their capacity to produce ATP, relative to their overall ATP consumption (Figure 4Ai-ii). The ETC-ind state could result from long-term changes in glycolytic capacity, from increased storage metabolites (such as glycogen), or from reduced anabolic demand. To discriminate between these possibilities, we cultured cells in the absence of glucose for 24 hours, and then added glucose for a short time window (1 minute or 30 minutes) prior to treating with oligomycin (Figure 4B,C). Because short exposure to glucose is unlikely to be sufficient to increase the expression of glycolytic machinery or storage pools, ETC-ind cells relying on these mechanisms would become ETC-dep during the starvation period (Figure 4Aiii). However, ETC-ind cells were detected at a much higher frequency than when cultured in glucose continuously. This result implies that, over the starvation period, ATP consumption rates decline more than glycolytic capacity, resulting in anabolically inactive cells that have relatively low demand, but remain poised to utilize glucose when it is resupplied (Fig. 4Aiv). At the same time, the frequency of ETC-ind cells remained dependent on the availability of glucose (Figure 4C), further supporting that the distribution between ETC-ind and ETC-dep states is defined by both anabolic ATP usage and glycolytic production.

**Figure 4.**
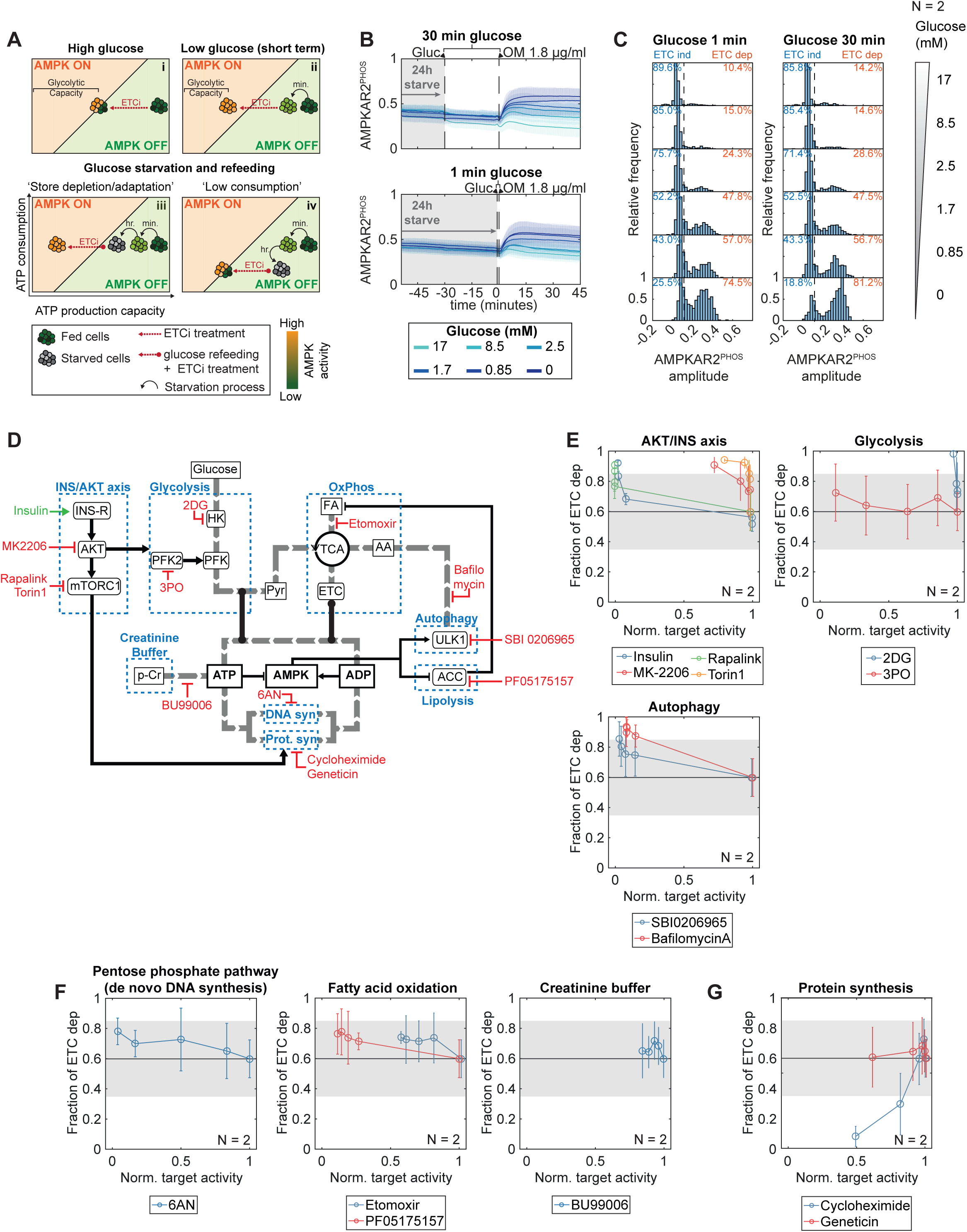
The balance between glycolysis and protein translation regulates AMPK heterogeneity. A: Cellular metabolic states as a function of ATP production (horizontal axes) relative to ATP consumption (vertical axes). The diagonal line in each panel represents the amount of ATP production capacity that is needed to meet a given level of ATP consumption without perturbing energy charge and triggering AMPK activity. Under high glucose (i), cells have excess ATP production capacity, but ETCi treatment reduces ATP production acutely. Some cells retain sufficient glycolytic capacity to meet the demands of consumption, averting AMPK activation. Under low glucose (ii), glycolytic capacity is immediately reduced and all cells fail to satisfy ATP demands following ETCi, and thus fall in the ETC-dep region. (iii-iv) Hypotheses for metabolic adaptation to long-term glucose starvation. In (iii), starvation may cause cells to deplete metabolite stores or may change their gene expression profile to reduce glycolytic capacity; upon re-supply of glucose followed immediately by ETCi treatment, such cells would score as ETC-dep. Alternatively (iv), glucose-starved cells may reduce their ATP consumption by downregulating energy-consuming processes; upon glucose resupply and ETCi treatment, these cells would need less ATP production capacity to meet their demands and would score as ETC-ind. B: Average AMPKAR2^PHOS^ measurements for MCF10A cells grown in the absence of glucose for 24 hours and then treated with glucose at specified concentration, followed by oligomycin 1.8 µg/ml at 1 minute or 30 minutes after glucose addition. C: Histogram of AMPKAR2^PHOS^ amplitudes after oligomycin treatment for the conditions shown in (B). D: Schematic of metabolic processes and regulation that potentially affect ATP production and consumption. Small molecule inhibitors targeting these pathways are shown in red. E, F, G: Redistribution of AMPKAR2^PHOS^ amplitude responses in response to inhibitor treatment. MCF10A cells were cultured in growth medium containing 17 mM glucose, and treated with a range of inhibitor concentrations for 30 minutes prior to oligomycin addition. The horizontal axes show inhibitor doses as normalized drug target activity according to pharmacokinetic information in previous literature (See Supplemental Figure S4 for normalization values and for absolute drug concentrations). The vertical axes indicate the fraction of cells scored as ETC-dep. The horizontal black line indicates the fraction of ETC-dep cells under control treatment (DMSO); points falling outside the gray region are considered significant by t-test.

To identify specific processes governing the balance between ATP generation and ATP consumption, we used a panel of inhibitors (Figure 4D). We tested each inhibitor by applying it for 30 minutes before treating with oligomycin and quantifying the distribution of AMPKAR and Perceval responses. An advantage of this approach is that, due to the short period of treatment, perturbations that shift the response distribution are expected to be limited to direct regulators, rather than indirect regulators (such as those that alter transcriptional profiles) that would be identified by longer-term knockdown approaches. To account for the fact that each inhibitor does not have equal efficiency on its target, we report changes in AMPK and Perceval response as a function of the expected residual target activity, based on published pharmacokinetic information for each inhibitor (Figure S4A).

Within the panel, inhibitors of the insulin/AKT signaling axis, glycolysis, or autophagy (Figures 4E and S4B) significantly shifted cells toward ETC-dep responses in a dose-dependent manner. Among the most sensitive targets were hexokinase (2-DG) and Akt (MK-2206), both essential for glucose catabolism, with significant effects even below target IC_50_. Torin1 (which inhibits both mTORC1 and mTORC2 complexes) was more effective than Rapalink (which is selective for mTORC1)(Rodrik-Outmezguine et al., 2016) in shifting cells toward ETC-dep; the higher sensitivity to Torin-1 is consistent with the role of mTORC2 in enhancing Akt activity. Similarly, the effect of inhibiting autophagy can potentially be explained by the finding that autophagy facilitates glucose transport (Roy et al., 2017). Inhibitors of other metabolic pathways showed lower sensitivity, including those targeting the pentose phosphate pathway (6AN), creatinine phosphate (BU99006), or fatty acid oxidation (etomoxir, PF05175157) (Figures 4F and S4B). In contrast to the increase in ETC-dep cells upon glycolysis blockade, inhibition of protein synthesis by cycloheximide (CHX) produced a strong decrease in ETC-dep responses (Figure 4G). This shift is consistent with the energy requirements for protein translation, which can account for 20% or more of ATP consumption within a cell (Buttgereit and Brand, 1995). Reducing this consumption would diminish the requirements for ATP production, thus lowering the impact on energy charge and AMPK activation upon ETCi treatment. Together, these results support that ETC-ind and ETC-dep cell states depend on a balance between the capacity to generate ATP through glycolysis and the rate of ATP usage by processes such as protein translation.

The insulin/Akt signaling pathway can potentially affect AMPK activation through multiple routes, including direct phosphorylation of AMPK (Suzuki et al., 2013), regulation of mTOR (which would increase protein translation and thereby ATP consumption), or control of glucose uptake (which could enhance the rate of ATP production by glycolysis). To determine whether ETC-ind cells can result solely from an increase in glucose uptake, we overexpressed the glucose transporter GLUT1. A co-translated mCherry fluorescent protein enabled us to quantify the degree of GLUT1 overexpression on a cell-by-cell basis. Independently of insulin treatment, higher expression levels of mCherry correlated with weaker AMPK responses (Figure 5A). In the absence of insulin, 60% of cells expressing GLUT1 were ETC-ind, while almost 100% of cells not overexpressing GLUT1 were ETC-dep, indicating that increased GLUT1 transporter alone is sufficient to confer ETC-ind behavior on cells.

**Figure 5.**
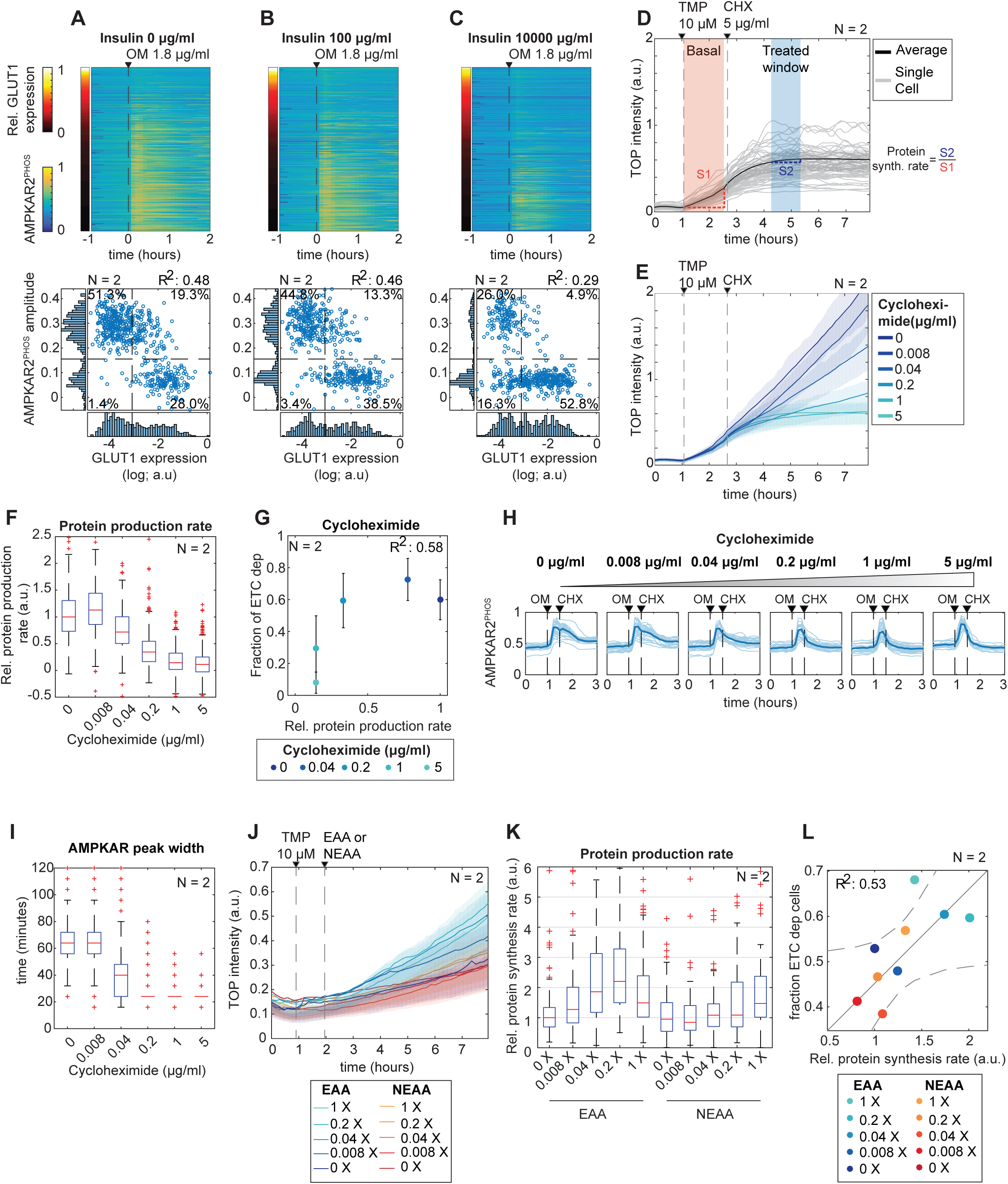
The balance between glycolysis and protein translation regulates AMPK heterogeneity. A, B, C: MCF10A-AMPKAR2 cells stably overexpressing GLUT1-IRES-NLS-mCherry were cultured with 0 µg/ml (A), 100 µg/ml (B) and 10000 µg/ml (C) insulin and exposed to oligomycin. Upper panels show heatmaps of AMPKAR2^PHOS^. Each row in the heatmap represents the time course of an individual cell; rows are sorted by relative mCherry intensity (which corresponds to the level of GLUT1 overexpression) represented by the color bar to the left of the red line. Lower panels show corresponding scatter plots of mCherry intensity and AMPKAR2^PHOS^ amplitude following oligomycin treatment. The percentage of cells falling within each quadrant is indicated. Quadrant boundaries (dashed lines) were determined by bimodal fitting across all samples. D: Measurement of protein translation rate in live cells. MCF10A cells stably expressing H2B-TOP-YFP-DD were treated with the degron inhibitor trimethoprim (TMP). Protein production rate was calculated as the slope of YFP intensity change during the 60 minutes after TMP treatment (orange shaded area). The effect of inhibitors on protein production were quantified as the slope for a 60 minute period beginning 90 minutes after treatment (blue shaded area). The translation inhibitor CHX is shown as a positive control. The relative protein production rate is calculated as the ratio of slopes in the blue and orange regions. E: Representative mean translation reporter results for a concentration series of CHX. Shaded areas show interquartile ranges. F: Boxplot of calculated single-cell relative protein production rates for the indicated dose of cycloheximide. Each box represents the distribution of >400 cells. G: Scatter plot showing popultion average relative protein production rate as measured by H2B-TOP-YFP-DD when treated with cycloheximide and their corresponding fraction of cells that become ETC dep after challenged with oligomcyin. H: Sample of AMPKAR2^PHOS^ when treated with oligomycin and later with cycloheximide at dose indicated. OM – oligomycin. Dark lines – AMPKAR2^PHOS^ population average. Light lines – single cell AMPKAR2^PHOS^ measurement I: Boxplot of single cell AMPKAR2^PHOS^ pulse widths after CHX treatment. Pulse widths were calculated as the time at which AMPKAR^PHOS^ decreased to 50% of the maximum value for each cell following treatment with CHX. J: Representative mean translation reporter results for MCF10A cells cultured in essential or non-essential amino acid at the indicated concentrations. Shaded areas shows the interquartile ranges. K: Boxplot of single-cell relative protein productions rate under the indicated amino acid conditions. Each box represents the distribution of >200 cells. L: Scatter plot of measured mean protein synthesis rates (I), and the fraction of ETC-dep cells for the corresponding amino acid concentrations.

We next examined if the shift away from ETC dependence induced by CHX can be attributed to regulation of protein synthesis, rather than a systemic stress response. A live-cell protein translation reporter, H2B-TOP-YFP-DD (Han et al., 2014b) was used to quantify cellular protein synthesis rates under different conditions, by calculating the slope of fluorescence increase following the addition of the stabilizer trimethoprim (Figure 5D). Using this readout, we were able to more directly relate protein synthesis rates to ETCi responses within MCF10A cells. Over a concentration curve of CHX (Figures 5E-F), ETC-dep cells decreased sharply at doses in which protein translation was inhibited by more than 60% (Figure 5G). Geneticin was unable to inhibit translation by more than 50%, even at high doses, accounting for its lack of an effect on ETCi responses (Figures S5A-C). Next, to observe the effects of protein synthesis inhibition even more acutely, we treated cells with CHX or geneticin 15 minutes after an AMPK response was initiated by oligomycin. Both inhibitors were sufficient to immediately halt AMPK activity, shortening the AMPK pulse length from an average of 60 minutes to 20 minutes (Figures 5H-I and S5D-E). To verify that the protein translation rate exerts an effect independently of chemical inhibitors, we cultured cells in the presence of varying extracellular concentrations of amino acids. Variation in essential amino acids (EAA), and to lesser degree non-essential amino acids (NEAA), modulated the rate of protein synthesis as measured by the translation reporter (Figures 5J-5K). Cells cultured in these conditions were then treated with oligomycin, and AMPK responses were measured, demonstrating a correlation between protein synthesis rate and the fraction of ETC-dep cells (R^2^ =0.53, Figure 5L). These results support the concept that ETC-ind cells depend on a low rate of protein synthesis to enable ATP production by glycolysis to maintain cellular energy charge above the threshold to trigger AMPK.

To establish whether heterogeneity in AMPK responses controlled by insulin is a common feature shared by other cell types, we stably expressed AMPKAR2 in a series of other cell lines, including 184A1 (immortalized mammary epithelial), MCF7 (hormone receptor-positive breast cancer), U87 (glioblastoma), A549 (non-small cell lung cancer, LKB1-deficient). These cell lines were cultured in varying concentrations of insulin, and challenged with oligomycin. Similar to MCF10A, 184A1 cells responded with a bimodal distribution of AMPK responses that shifted upon insulin treatment to favor the ETC-ind population (Figures S5F-5G). However, the frequencies of ETC-ind and ETC-dep cells varied relative to MCF10A. Nearly 100% of 184A1 cells scored as ETC-ind in the presence of insulin, which fell to 66% ETC-ind in the absence of insulin, suggesting that these cells have, on average, a higher capacity to meet their ATP requirements through glycolysis alone. MCF7 cells displayed a configuration shifted in the opposite direction: 100% of cells showed a robust AMPK response to oligomycin in the absence of insulin, and were only moderately shifted toward ETC-ind by insulin. These distributions can be explained as a higher dependence on OxPhos in this cell type, either due to a lower capacity for glycolysis, or higher cellular demands. U87 cells also showed a broad distribution of responses, which shifted toward ETC-ind with insulin treatment. A549 cells, which are deficient for the essential AMPK activator LKB1, showed very little AMPK response regardless of insulin treatment. Overall, these responses indicate that heterogeneity in cellular energetic configuration is widespread among human cell lines.

### Variation in AMPK activity is strongly coupled to the growth regulatory network

AMPK is known to inhibit the activity of both mTORC1 (Gwinn et al., 2008; Inoki et al., 2003) and the Ras/ERK pathway (Shen et al., 2013) (Figure 6A), which promote cell growth and proliferation, respectively. We investigated whether the magnitude of AMPK response mounted by each cell impacts the activity of these connected signaling pathways. To monitor ERK activity simultaneously with AMPK activity, we used a translocation-based reporter, ERKTR (Regot et al., 2014) (Figure S6A). Upon oligomycin treatment, ERKTR detected, on average, a decrease in ERK activity (Figure S6C), consistent with inhibition of this pathway by active AMPK. When considered on a cell-by-cell basis, the reduction of ERK activity correlated with the magnitude of AMPK activation (Figure 6B), whereas no correlation was found in the absence of oligomycin. Furthermore, when time courses of AMPKAR2 and ERKTR signals were tracked in individual cells, a clear anti-correlation was observed over the course of more than 12 hours. Over this time, pulses of AMPK activity were matched by depressions in ERK activity (Figures 6C and S6C,E), with a lag time of 6 minutes or less (Figure 6E). The Pearson correlation coefficient between the aligned time series indicated a strong statistical significance (Figure 6F).

**Figure 6.**
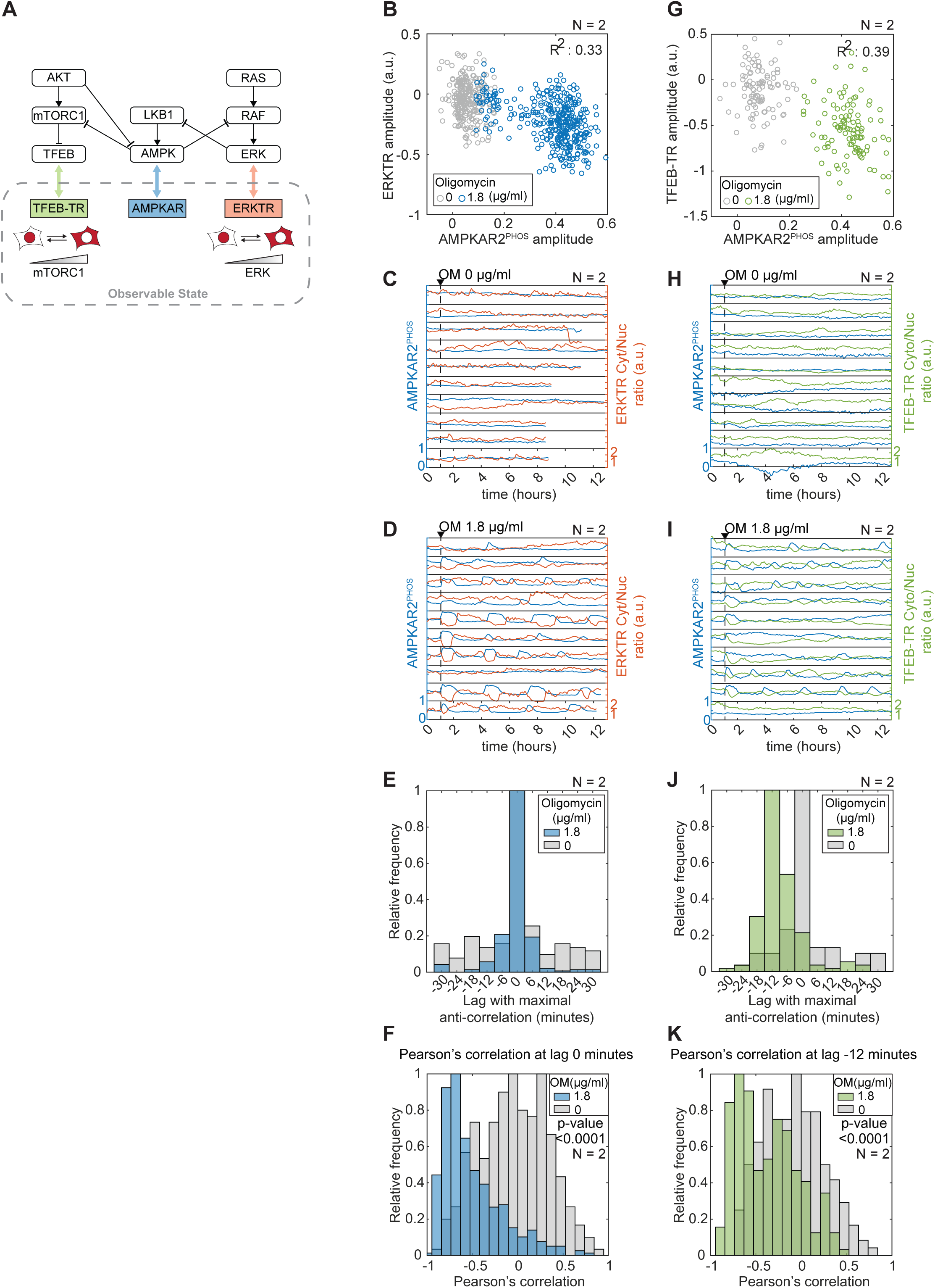
Heterogeneous AMPK responses correlate with perturbation of downstream signalling. A: Schematic of connections between AMPK, mTORC1 and ERK, and corresponding reporters for live-cell analysis. B: Scatter plot of single-cell AMPKAR2^PHOS^ amplitude and corresponding ERKTR changes in cells treated with oligomycin or vehicle (DMSO). C, D: Representative single-cell profiles of AMKPAR2^PHOS^ and ERKTR cytoplasmic-to-nuclear ratio for cells treated with oligomycin 1.8 µg/ml (C) or vehicle (D). Each panel shows one cell. E: Lag analysis of cross-correlation between AMPKAR2 and ERKTR using adaptive dissimilarity index, showing maximal correlation at 0 minute-lag F: Histogram of cross-correlation coefficients between AMPKAR2 and ERKTR signals at 0 minute lag when treated with oligomycin (blue) or vehicle (gray). G: Scatter plot of single-cell AMPKAR2^PHOS^ amplitude and corresponding TFEBTR changes in cells treated with oligomycin or vehicle H, I: Representative single-cell profiles of AMKPAR2^PHOS^ and ERKTR cytoplasmic-to-nuclear ratio for cells treated with 1.8 µg/ml oligomycin (C) or vehicle (D). J: Lag analysis of cross-correlation between AMPKAR2 and TFEBTR using adaptive dissimilarity index, showing maximal correlation at −12 minute-lag K: Histogram of cross-correlation coefficients between AMPKAR2 and TFEBTR signals at −12 minute lag when treated with oligomycin (green) or vehicle (gray).

To detect mTORC1 activity in live cells, we used the nuclear-to-cytosolic translocation of a fluorescent protein fusion to transcription factor EB (TFEB-TR), which is stimulated by direct mTORC1-mediated phosphorylation (Figure S6B) (Li et al., 2018; Settembre et al., 2012). As in the case of ERKTR, TFEB nuclear-to-cytosolic ratio was decreased following oligomycin treatment (Figure S6D) and correlated to AMPKAR2^PHOS^ at the single cell level (Figure 6G). Cycles of TFEB translocation coincided with AMPK pulses, following a ∼12 minute lag (Figures 6H-6K and S6F). While we cannot rule out additional mechanisms of activation, such as phosphorylation of TFEB by other kinases, these results are consistent with dynamic regulation of mTORC1 by AMPK. Moreover, consistent with a change in mTORC1 activity, we also observed suppression of protein translation as indicated by the H2B-TOP-YFP-DD (Figure S6G-H). Overall, these results support a model in which variation in AMPK activity determines the extent of inhibition of ERK and mTORC1 kinase activity during energy stress.

## Discussion

Our results establish a kinetic model of metabolic variation within individual cells, both under metabolic stress (ETCi) and during normal growth in culture. The variation we observe can be conceptualized as an “inner” cycle in which periodic switching of AMPK activity is enforced by ETCi, and an “outer” cycle that fluctuates spontaneously in growing cells without AMPK activity (Figure 7). Our data imply that the underlying cellular metabolic configuration – that is, the balance of glycolytic capacity relative to ATP demand-shifts over time as cells progress through the outer cycle. The state of this outer cycle at the time of ETCi treatment determines the point at which each cell enters the inner cycle (Figure 7, red arrows), and thus defines the initial response to ETCi treatment. Once forced into the inner cycle by ETCi, the balance between glycolysis and ATP demand becomes tightly coupled to ADP/ATP ratio, enabling them to be observed by the Perceval reporter or AMPK activity. Unlike the inner cycle, the progression of the outer cycle is not directly detectable by AMPK activity measurements, likely because when both glycolysis and OxPhos are active, ADP/ATP ratio can be maintained more easily, masking fluctuations in either one of the two processes. However, the initial response of AMPK or ADP/ATP ratio upon ETCi treatment provides an observable that can be used to reveal the distribution of underlying metabolic states. Because this observation is indirect, it remains difficult to characterize the periodicity of the outer cycle; although it could be similar to the 3-5 hour period of the inner cycle, it is not clear to what extent ETCi treatment perturbs the free-running outer cycle. An important next step will be to identify more expedient observables to monitor the state of the outer cycle.

**Figure 7.**
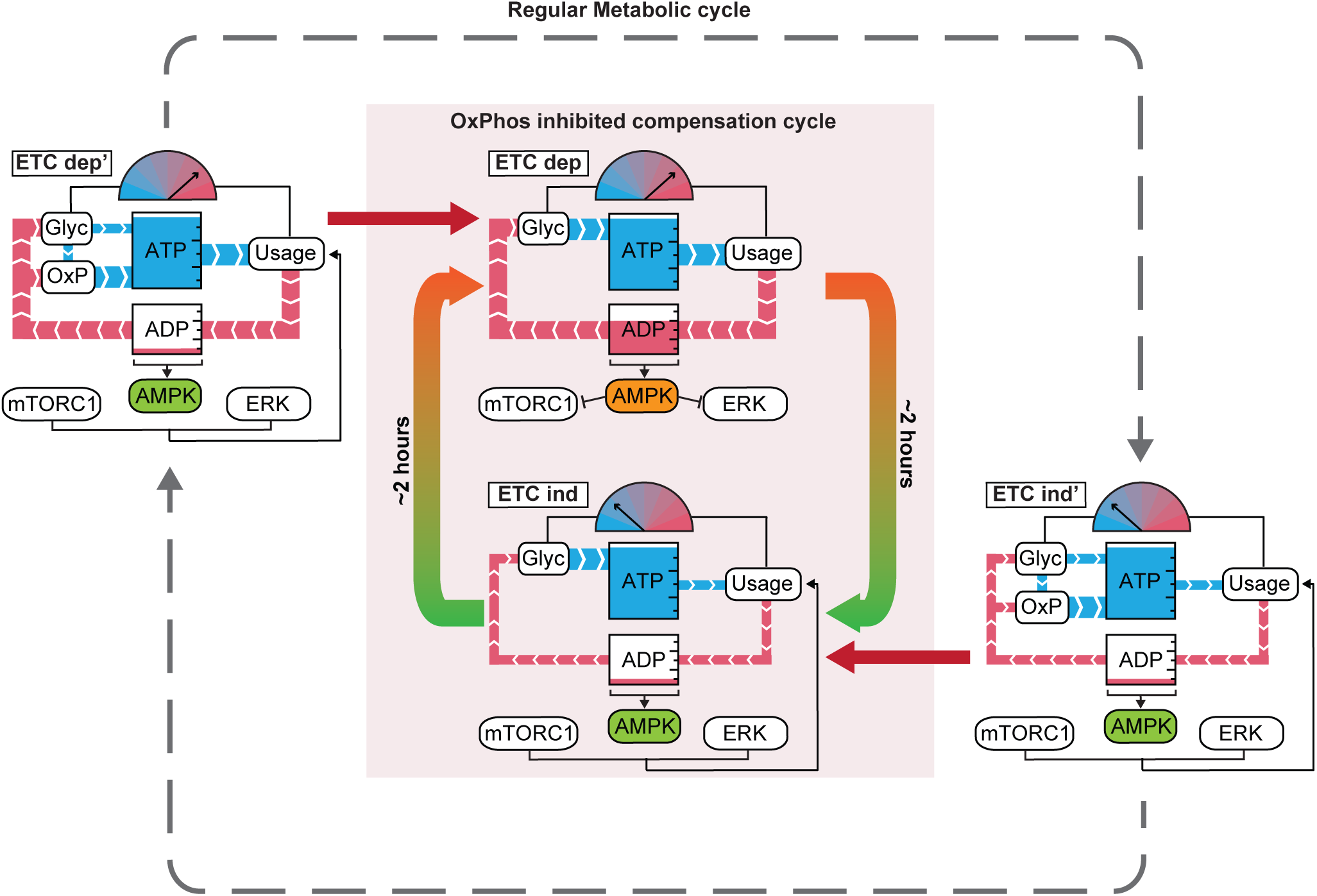
A model of metabolic cycling in mammalian cells. Simplified diagrams are used to indicate the state of central ATP metabolism at different points in time. Blue “pipes” indicate flux of ATP, and red pipes the flux of ADP. Meter icons indicate the balance of glycolytic ATP production capacity relative to ATP consumption, with blue indicating excess production capacity, and red indicating excess consumption. The dotted inner border indicates the dynamics of the network in the presence of ETCi, in which OxPhos is inactivated and the state of the network (i.e. the meter) is directly coupled to AMPK activity. The dynamics of switching between these states can be clearly observed (e.g., Fig. 1D). The outer cycle indicates the inferred dynamics of metabolic balance in the absence of ETCi. The dynamics of this cycle can be observed indirectly by monitoring whether, upon ETCi treatment, a cell enters the inner cycle in the ETC-dep state (A → A’) or in the ETC-ind state (B → B’). Because OxPhos is active for cells in the outer cycle, AMPK activity remains low regardless of whether the energy balance favors ATP consumption (A) or glycolysis (B).

In mammalian cells, cycling of metabolic parameters has primarily been linked to the cell division cycle (CDC) (Ahn et al., 2017) or to circadian rhythms (Bass and Takahashi, 2010). In yeast, a more extensive system of metabolic oscillations has been identified (Tu et al., 2005), termed the yeast metabolic cycle (YMC). The YMC is often coupled to the cell cycle during rapid growth (Tu et al., 2007), but can also progress independently of the CDC (Papagiannakis et al., 2017; Silverman et al., 2010), and can modulate cell cycle progression (Argüello-Miranda et al., 2018; Papagiannakis et al., 2017; Shi and Tu, 2013). The existence of this complex system in yeast suggests that more extensive metabolic oscillations also exist in mammalian cells but have not been observed. However, it is likely that the observables of the YMC will not be directly transferrable; for example, fluctuations in ATP concentration can be tracked using Ateam in yeast (Papagiannakis et al., 2017) but appear to be more effectively buffered in mammalian cells. The cycles we identify here suggest alternation between at least two phases: one in which protein synthesis and anabolic processes consume ATP at a low rate that can be sufficiently supplied by glycolysis, and one in which the anabolic activities of the cell require OxPhos to maintain energy charge. These phases show similarity to the YMC in both their approximate period (3-5 hours in the case of the inner cycle) and in the alternation of cellular dependence on OxPhos. The ETC-dep phase we observe may be equivalent to the “OX” phase of the YMC, in which protein translation and other anabolic processes accelerate, fueled by a higher rate of OxPhos and oxygen consumption (Cai and Tu, 2012). The behavior of ETC-ind cells is consistent with certain characteristics of the “reductive/charging (RC)” phase of the YMC, including its association with cellular quiescence and autophagy (Slavov et al., 2011). While we identify a number of conditions capable of forcing cells from one state to another, the question of what mechanisms drive this transition under free-running conditions remains an open challenge. The possibility that there is an underlying cycle between translation and autophagy, orchestrated by opposing control of ULK1 by mTOR and AMPK (Szymańska et al., 2015), is largely consistent with our data, but still speculative.

A remarkable aspect of metabolic kinetics apparent in our analysis is the stability of ATP concentration even under severe perturbation of ATP production. Our analysis of glucose and oxygen consumption under both uninhibited and ETC-inhibited states implies that a large fraction of ATP production can be shifted to glycolysis within seconds, despite its low yield of 2 ATP per glucose molecule relative to the ∼30 produced by OxPhos. Because ATP homeostasis is maintained even in cells without a detectable AMPK response (ETC-ind), AMPK is likely not required for this initial adaptation of ATP flux to ETCi. Moreover, two of the main routes of AMPK-mediated energetic balance - glucose uptake and protein translation rate – are only modestly altered upon ETCi treatment (Supplemental Figures S1 and S6). Rather, our data imply that during ETCi treatment, flux through glycolysis is redirected from the production of biosynthetic intermediates, which are uncoupled from ATP production (Lunt and Vander Heiden, 2011), to prioritize the production of ATP. In ETC-ind cells, the current anabolic load is low enough that this shift can occur without a large perturbation of ADP/ATP ratio, resulting in no detectable activation of AMPK. In ETC-dep cells, this shift is still rapid enough to preserve ATP levels, but generates a large enough rise in ADP/ATP ratio to provoke high AMPK activity. The observation that ETC-ind cells eventually (and repeatedly) enter the ETC-dep phase during prolonged ETCi treatment indicates that this cycle of anabolic activity persists despite the reconfiguration of glycolysis.

Our results clarify the temporal role of AMPK in metabolic adaptation. AMPK activity is essential for mammalian viability (Viollet and Foretz, 2016) and is often termed a “master regulator” of cellular metabolism (Witczak et al., 2008). However, AMPK is not required for *cellular* viability, even under metabolic stress (O’Neill et al., 2011), suggesting that it is not the primary mechanism by which cells acutely adapt to energetic challenge. Our results are consistent with this lack of an immediate role, as AMPK activity is not homogenous or rapid enough to explain the resistance of ATP levels to ETCi. Instead, the dynamics of AMPK activation indicate that it detects the shift in ADP and AMP that results from rapid buffering during the initial response, and relays this information to regulate its downstream targets, including mTORC1 and ERK kinase activity. Other metabolic effects of AMPK, such as phosphorylation of ACC, occur on a similar timescale (Jeon et al., 2012). Our results solidify and extend a model in which there are multiple distinct timescales involved in energetic homeostasis (Hardie et al., 2012): rapid buffering at the level of metabolite flux (seconds), which triggers AMPK-mediated adaptive changes (minutes-hours), both of which are subject to the independent underlying cycle of anabolic activity that we identify here (>4 hours).

Activation of AMPK and its downstream effectors may be beneficial for treating diabetes, cancer, and other diseases, and there have been a number of efforts to develop pharmacological AMPK activators (Cokorinos et al., 2017; Myers et al., 2017). In particular, biguanides are an important class of diabetes drug that act in part as ETC inhibitors, and which induce cyclic AMPK activation similar to that shown here (Hung et al., 2017a). Understanding the kinetics of AMPK in response to ETCi challenges, and the factors that underlie the heterogeneous ETCi response, will be important in optimizing AMPK activation strategies for beneficial cellular effects. The ability to predict and control the fraction of cells that respond to ETCi may allow these drugs to be tailored toward different goals. Potent induction of energy stress in the largest number of cells possible may be desirable in the case of anti-cancer therapy, but intermittent activation may be preferable when trying to restore physiological energy balance in diabetes or metabolic syndrome. It may also be the case that cellular responses to other drugs, including cytotoxic chemotherapies and targeted kinase inhibitors, depend in part on the state of the underlying anabolic/catabolic cycle, because both cell death and proliferative signaling are tightly interconnected with cellular metabolism.

## Acknowledgements

Funding for this work was provided by the American Cancer Society (IRG-95-125-13), the National Institute of General Medical Sciences (1R01GM115650), and the American Association for Cancer Research/Stand Up To Cancer (SU2C-AACR-IRG-01-16). Stand Up To Cancer is a program of the Entertainment Industry Foundation. Research grants are administered by the American Association for Cancer Research, the scientific partner of SU2C. Flow-cytometry services were supported by the UC Davis Comprehensive Cancer Center Support Grant (CCSG) awarded by the National Cancer Institute (NCI P30CA093373). Metabolomics study was performed in collaboration with West Coast Metabolomic Center [WCMC]. Seahorse assay was performed in collaboration with Cortopassi lab, UC Davis - school of veterinary medicine. We thank Marek Kochanczyk, Ben Tu, Ralph DeBerardinis, and Joshua Rabinowitz for helpful discussions.

## Abbreviations

OM: oligomycin
INS: insulin
ETC: Electron Transport Chain
OxPhos: Oxidative Phosphorylation
ETCi: Electron Transport Chain inhibitor
ERKTR: ERK kinase translocation reporter
TFEBTR: TFEB translocation reporter

## Materials and methods

### Reporter construction

AMPKAR2 (Hung et al., 2017b) and ERKTR-mCherry (Sparta et al., 2015)were previously described. PercevalHR (Tantama et al., 2013b), ATeam1.03 (Imamura et al., 2009a), H2B-TOP-YFP-DD (Han et al., 2014a), and GLUT1 were obtained from Addgene. PercevalHR was modified with a nuclear export sequence at the C-terminus to direct it to the cytosolic compartment. AMPKAR2, PercevalHR, and Ateam1.03 sensors were expressed in pPBJ, piggyBAC transposase-mediated delivery systems (Yusa et al., 2011) to minimize recombination between CFP and YFP. GLUT1-IRES-NLS-mCherry was constructed by cloning the GLUT1 coding sequence (Takanaga et al., 2008) into the retroviral vector pBabe-neo (BamHI/XhoI); IRES-NLS-mCherry was then inserted at the 3’ end of GLUT1 (XhoI/SalI). TFEBTR-mCherry was constructed by inserting the coding sequence for TFEB residues 1-247 into pLJM1 upstream of and in-frame with the coding sequence of mCherry. were expressed in lentiviral expression system using, pLJM vector. Correct insertions for all plasmids were confirmed by sequencing.

### Reporter Delivery

Cell lines stably expressing biosensors were generated by retroviral transduction or transfection with the PiggyBac transposase system (Yusa et al., 2011). All PiggyBac plasmids were delivered by electroporation (Amaxa II system, Lonza). After transfection or transduction, cells were selected with puromycin (1–2 μg/ml) or geneticin (300 μg/ml); single-cell clones were made by limiting dilution or flow cytometry sorting. For each reporter, we isolated multiple stable clones with homogenous expression; all data reported in this study reflect representative behaviors that were consistent across all clones of each reporter (a minimum of three clones in each case). All reporter lines were confirmed to be mycoplasma-negative by PCR; results were validated by third-party testing of selected lines (ATCC).

### Cell culture and media

Routine cell culture for human mammary epithelial MCF10A cells clone 5E (Janes et al., 2010) and 184A cells were performed as previously described (Debnath et al., 2003). MCF10A and 184A1 were grown in ‘DMEM/F12 growth medium’ (see Media table). Primary stocks from the original clonal derivation (MCF10A-5E) or from the ATCC (184A1) were used in all experiments. MCF7, U87, and A549 cell lines were obtained from ATCC and cultured in ‘DMEM growth media’(see Media Table). All cells were routinely split when they are ∼80% confluent.

In live microscopy experiments, we used a custom formulation, termed ‘imaging base-DMEM/F12’, which consists of DMEM/F12 lacking glucose, glutamine, riboflavin, folic acid, and phenol red (Life Technologies or UC Davis Veterinary Medicine Biological Media Service) to avoid fluorescence background. All experiments involving MCF10A or 184A1 cell line were performed in ‘Imaging medium 1’ (see Media table). For experiments with MCF7, U87 or A549 cell lines, ‘Imaging medium 2’ (see Media table) was used. For all experiments, ‘Imaging medium 1’ and ‘Imaging medium 2’ were supplied with glucose 17 mM and 25 mM, respectively, unless indicated otherwise.

Before imaging, cells were washed twice with their respective media and then cultured in imaging experiment media at least 2 hours prior to imaging, unless indicated otherwise. The cell to media ratio was maintained at 150-200 cells/µl for all experiments. For experiments involving titration of insulin or EGF concentrations, cells were placed in EGF- or insulin-deficient media for 4 – 6 hours prior to imaging.

### Live-cell fluorescence microscopy

Time-lapse wide-field microscopy was performed as described previously (Hung et al., 2017b; Pargett et al., 2017). Briefly, 25000 cells were seeded one day prior to imaging in glass-bottom 96-well plates (Cellvis P96-1.5H-N, Mountain View, CA) pretreated with type I collagen (Gibco A10483-01) to promote cell adherence. For experiments with drug addition, cells were placed in imaging medium until the addition of drug. For drugs dissolved in DMSO, the final DMSO concentration was <0.1%. Cells were maintained in 95% air and 5% CO_2_ at 37°C in an environmental chamber. Images were collected with a Nikon (Tokyo, Japan) 20/0.75 NA Plan Apo objective on a Nikon Eclipse Ti inverted microscope, equipped with a Lumencor SOLA or Lumencor SPECTRA X light engine (see Lightsource table). Fluorescence filters used in the experiment are: DAPI (custom ET395/25x - ET460/50m - T425lpxr, Chroma), CFP (49001, Chroma), Sapphire (custom ET420/10x - ET525/50m - T425lpxr, Chroma), GFP (49002; Chroma), YFP (49003, Chroma), Cherry (41043, Chroma) and Cy5 (49006, Chroma). For AMPKAR2 and Ateam1.03 biosensor CFP and YFP filters were used to acquired image, while PercevalHR biosensor Sapphire and GFP filters were used. Images were acquired using Andor Zyla 5.5 scMOS camera every 6 – 7 minutes with at 2×2 binning. Exposure time for each channel are 25-50 ms for DAPI; 150 – 250 ms for CFP; 150 – 250 ms for YFP; 500 – 750 ms for Sapphire; 500 – 750 ms for GFP; 300 – 500 ms for Cherry and 300 – 500 ms for Cy5.

### Immunofluorescence microscopy

Cells were plated in glass-bottom plates as described above. At indicated times during live-cell imaging experiment, 8% paraformadehyde was added directly into imaging media to make 2% paraformaldehyde final concentration. Paraformaldehyde fixation was performed for 15 minutes, followed by permeabilization with 100% methanol. Cells were then washed in PBS-T (0.1% Tween-20 in PBS) twice and blocked with Odyssey Blocking Buffer (Li-Cor, Lincoln, NE) for 1 hour at room temperature. Samples were then incubated with primary antibody against Phospho-Acetyl-CoA Carboxylase (Ser79) (Cell Signalling 11818; dilution 1:1000) diluted in blocking buffer overnight at 4 °C. Secondary staining was performed with Alexa647-conjugated anti-rabbit (Life Technologies, A-21245, diluted at 1:1000 in blocking buffer), followed by DNA staining with Hoechst-33342 (Life Technologies, H3570, diluted at 1:1000 in PBS). Plates were imaged as described for live-cell microscopy, using DAPI and Cy5 filter sets.

### Image processing

After background subtraction and flat field correction, imaging data were processed to segment and average pixels within each identified cell’s nucleus and cytoplasm, using a custom procedure written for MATLAB (Pargett et al., 2017), with modifications in the cytosolic identification protocol as described below. Image data were stored in ND2 files generated by NIS Elements and accessed using the Bio-Formats MATLAB toolbox. Individual cells were tracked over time using uTrack 2.0 (Jaqaman et al., 2008). Cytoplasmic masks were created by watershed method (Vincent and Soille, 1991) using cytosolic YFP (for cell lines expressing AMPKAR2 or ATeam1.03) or GFP (for cell lines expressing PercevalHR) to identify the cytosolic boundary. The cytosolic area is further restricted to the area within 5 pixels from the nuclear border. The resulting single-cell time series traces were filtered for quality by minimum length of trace and maximum number of contiguous missing or corrupt data points.

### FRET reporter measurement

To quantify FRET biosensors (AMPKAR2 and ATeam1.03), we employed a spectral model of light propagated through the microscopy system, including the live cell specimen. In our microscope, light is generated via LEDs and passed through spectral excitation filters to illuminate the specimen; fluorescent light from the specimen is collected by the objective and passed through spectral emission filters before illuminating the camera. We therefore defined a spectral model including all elements that modify the spectrum: (1) light source, (2) excitation filters, (3) fluorophores in the specimen, (4) emission filters, and (5) camera sensor. For a given pixel, the model is the product of: light source intensity *I_LS_*(*λ*), excitation filter transmissivity *F_ex_*(*λ*), specimen absorbance *A_S_*(*λ*), specimen concentration *C_S_*, specimen quantum yield *QY_S_*, specimen emission spectrum *ϕ_S_*(*λ*), geometric fraction of light collected by the objective *G_geo_*, emission filter transmissivity *F_em_*(*λ*), camera quantum efficiency *QE_cam_*(*λ*), loss to absorbance/scattering from all optics *G_opt_*, and time of exposure *t*. All terms that do not have a uniform effect across the spectrum are shown as dependent on wavelength *λ*.

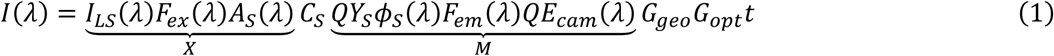

The linearity in absorption by the specimen implies the assumption that the concentration of specimen is low enough for the exponential in the Beer-Lambert law (transmissivity, *T*(*λ*)=10*^−A(λ)c^*) to be approximately linear. The model may be simplified slightly by lumping all uncertain gain terms (geometric collection *G_geo_*, optical loss *G_opt_*, light source average power, electronic gain in camera circuitry, any other inefficiencies, etc.) to a single term G = *G_geo_G_opt_G_ls_G_cam…._* All terms must be determined to fully calibrate the model and predict, for example, the concentration of fluorophore in the specimen.

Including the effect of a FRET interaction between two fluorophores in this model gives a general model, suitable for any spectral collection channels (Hoppe et al., 2002). This equation now includes in the integration over the spectrum, yielding the measured pixel intensity. Though not explicitly shown in notation from here on, all fluorophore-specific terms (*X^D^*, *M^A^*, etc.) are dependent on wavelength. Superscripts D and A refer to the Donor and Acceptor molecules, respectively. Note that while the FRET reporters used in this report are single molecules with two fluorophores, the concentration of the donor (*C^D^*) and acceptor (*C^A^*) can be considered separately, accounting for truncated proteins and bleached or denatured fluorophores, which may vary from cell to cell and across cultures.

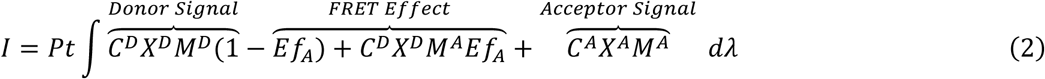

Our measurement of FRET sensors involves recording two spectral channels, CFP (donor fluorophore) and YFP (acceptor fluorophore), one of which (YFP) essentially accounts for the expression level of the sensor. By considering a ratio of these channels, gain terms that are common to both channels are eliminated, leaving the spectral terms, as well as exposure time and average light source power if these varied between channels.

The ratio of CFP to YFP is then represented by the following.

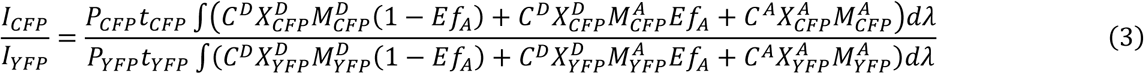

Because the CFP and YFP filter sets we employ are spaced spectrally such that CFP and YFP are not cross-excited, 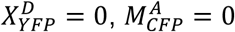, simplifying the model.

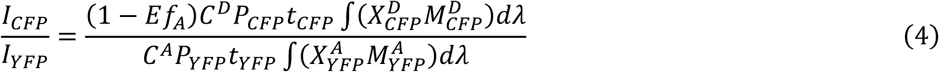

Solving for the fraction associated provides the necessary correction term, which can be considered as two factors: *R_C_*, the ratio of active Donors to Acceptors, which may vary from cell to cell due to bleaching and translation defects, and *R_P_*, the ratio of imaging power per channel, which is a constant per experiment. The FRET efficiency for both AMPKAR2 and ATeam1.03 (*E*) is not known, but has a strictly linear effect, leaving relative fractions associated accurate but absolute values uncertain.

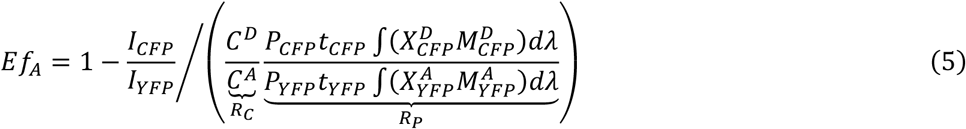

*R_P_* may be measured by careful calibration. We employ manufacturer specifications for all filter properties, camera spectral sensitivity, and light source spectra. We draw fluorophore properties and spectra from published literature. Finally, we calibrate for variation in light source and filter properties by measuring the excitation power transmitted through our light path to the specimen using an optical power meter and a photodiode sensor (ThorLabs PM100D and S120VC, respectively). However, *R_C_* remains uncertain and dependent on experimental conditions, and in this system contributes to technical error (and herein is given an assumed value of 1). Note that an additional bias factor from excess free Donor has been neglected this form as it is expected to be minor and is impractical to calibrate (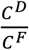 would be multiplied by the entire right-hand expression, *C^F^* being the true concentration of working FRET pairs).

### Perceval reporter measurement

Unlike FRET reporters, PercevalHR has only one fluorophore, cp173 mVenus, that binds to ATP and ADP differentially, resulting in a shift of excitation spectra with peaks at 470 nM (ATP-bound) and 405 nM (ADP-bound) (Tantama et al., 2013b). To measure the proportions of these forms, we imaged cells expressing PercevalHR reporter with Sapphire and GFP filters (see Live-cell fluorescence microscopy). To account for variation in microscope light source set up from experiment to experiment, we considered equation (1) and scale image measurements by the relative excitation intensity delivered in each channel – eq 6.

[*C^P^* – PercevalHR reporter concentration; *P_Sap_* and *P_GFP_* – fraction of ADP-bound and ATP bound PercevalHR, respectively; *t_Sap_* and *t_GFP_* – exposure time of Sapphire and GFP channels, repectively; *I_LS_*(*λ*) – light source intensity; *F_exSap_* and *F_exGFP_* - excitation filter transmissivity of Sapphire and GFP filters, respectively]

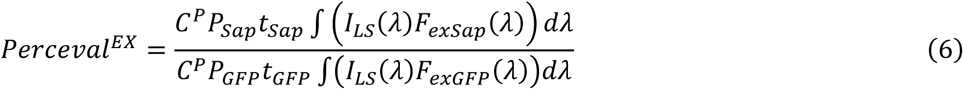

### AMPK substrate phosphorylation estimate

Since AMPKAR2 reporter is a substrate for AMPK kinase activity, it is possible to estimate the fraction of sensor that is phosphorylated using Phos-Tag™ electrophoresis, followed by immunoblot against GFP (see Phos-Tag electrophoresis and western blot). This measurement allows us to convert units of FRET fraction associated into the fraction of sensor that is phosphorylated, AMPKAR2^PHOS^, which is more meaningful. To achieve this goal, we perform estimate fraction of AMPKAR2 phosphorylated on average population responses via Phos-Tag western blotting for the phosphorylated form of the AMPKAR2 sensor. Lysates from the MCF-10A cell lines treated with condition indicated in Supplementary Figure 1B were used (4 replicates per cell line and treatment). These conditions were selected because they exhibit sustained AMPKAR2 activity, unlike AMPKAR2 response to ETCi. After Phos-Tag western blotting, membranes were stained with an anti-GFP antibody (Cell Signaling Technologies #2955) to visualize the AMPKAR2 reporter, and the average fraction of reporter phosphorylated was quantified. These values were then compared with the average fraction associated as calculated from live-cell experiments at corresponding treatments and time points. Linear fitting was performed and providing a calibrated measurement of the fraction of AMPKAR2 phosphorylated, based on live-cell measurements – eq 7.

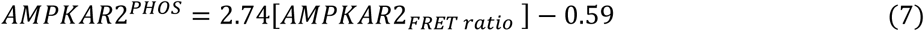

### Cell age and sister cell analysis

For sister cell analysis, we expressed an NLS-mCherry nuclear marker in MCF10A-AMPKAR2 cell line to improve nuclei tracking accuracy across cytokinesis. Cell division events were first automatically identified by uTrack2.0 (Jaqaman et al., 2008), and later manually verified. In total, we were able to record more than 5,500 cell division events (11,000 related cells) within 25 hours. This dataset gave us estimates on both each cell’s age and their lineage at the time they were challenged with oligomycin.

The similarity of AMPKAR2^PHOS^ response between cell sister cells was calculated by computing the Euclidean distance of AMPKAR2^PHOS^ responses within the 2-hour window after oligomycin treatment. To ask whether AMPKAR2^PHOS^ response between sister cells was more similar than that of unrelated cells, we generated 1000 random pairs of cells that divided at the same time and computed the average AMPKAR2^PHOS^ Euclidean distance. We were able to estimate Euclidean distance of AMPKAR2^PHOS^ between unrelated cells with 95% confidence interval. The age-dependent increase in the AMPKAR2^PHOS^ Euclidean distance was fitted by an exponential function to estimate the half-life.

### Drug Target Normalization

In order to account for the fact that the inhibitors shown in Figure 4D do not have equal efficiency on its target, we normalized the dose of each inhibitor by its target activity based on published reports (Figure S4A). For each drug, a one-term Hill function was fit using published dose curve values and used to predict residual target activity at each drug concentration used.

### Analysis and statistics of kinetics in reporter signals

A custom MATLAB algorithm was designed to identify peaks (Gillies et al., 2017) in the time-lapse signal of AMPKAR2, PercevalHR, and Ateam1.03 activity. The AMPK, PercevalHR, and Ateam1.03 were first smoothened using Butterworth low pass filter with 3-timepoints cut off period to remove spurious noise. Peaks and associated valleys in the index were identified by setting two local cutoff values, based on maximum and minimum values of the data within a sliding time window (typically 120 minutes for AMPKAR2 and PercevalHR, 30 minutes for Ateam1.03). A peak was detected if both cutoff values were crossed by a rise and subsequent fall in the index. Typically, more than 300 individual cell recordings were scored for each condition and plotted as a histogram.

### Bimodality test

To test for bimodality, data were fitted to a bimodal Gaussian mixture distribution and a panel of unimodal distribtions, including (including normal, log-normal, generalized extreme value, and Weibull). The best-fitted distribution was selected using corrected Akaike’s Information Criterion, to account for extra parameter terms (Cavanaugh, 1997). Data were considered bimodally distributed if and only if the bimodal Gaussian mixture distribution was ranked as the best-fitted distribution.

### Pearson’s cross-correlation of time series

The time series to be compared were normalized by subtracting by their corresponding averages. To quantify lag between reporters for each time series, the maximal cross-correlation value was computed using the MATLAB *xcorr* function. We assumed that each pair of reporters, namely AMPKAR2-TFEB-TR or AMPKAR2-ERKTR, has a characteristic lag time, estimated as the mode of calculated lags across all sampled cells. The lag identified from this process were used to align two time series data. Pearson’s correlation coefficient was computed from these aligned time series.

### Measurement of mitochondrial stress responses and ATP flux from glycolysis/oxidative phosphorylation

XF24 cell culture plates and sensor cartridges (100867–100) were purchased from Seahorse Bioscience (North Billerica, MA). Cells were seeded in XF24 cell culture plates at a density determined by optimization experiments and incubated at 37 °C with 5% CO_2_ overnight in growth medium; even distribution of cells was verified visually. For the mitochondrial stress test, the growth medium was completely removed 24 hours after plating, and cells were washed twice with 1,000 ml of pre-warmed ‘DMEM/F12 experiment media 1’. 500 ml of assay medium was added to each well and cells were incubated in a 37°C incubator without CO_2_ for 1 hr to allow cell equilibration with assay medium. Oxygen consumption rates were measured with the XF24 analyzer under this basal condition followed by sequential addition of varied oligomycin concentration as indicated in Supplementary Figure 1F. For ATP fluxes from glycolysis and oxidative phosphorylation estimation, the data collected in the previous study (Hung et al., 2017b) was applied to formula previously described by Mookerjee et al (Mookerjee et al., 2017b).

### Glucose consumption rate estimation

Extracellular glucose consumption rate estimation in MCF10A cell line was measured using a glucose/glucose oxidase assay kit (Thermo A22189). Briefly, MCF10A cells were plated in 96 well assay plate (Corning 3610) at 25000 cells/well in MCF10A full growth media overnight for attachment. 10 minutes prior to the experiment, growth medium was removed and replaced by ‘DMEM/F12 experiment media 1’ supplemented with 17 mM glucose at 40 ul per well. After pre-incubation, cells were treated with inhibitors and growth factors as indicated in Supplementary Figure 1H for 3 hours before media was sampled for glucose measurement. Samples of media were mixed with glucose oxidase reagent per manual provided in the kit. Luminescence was monitored by a microplate reader (Molecular Device, SpectraMax M5) at 560 nM, room temperature. Glucose consumption rate was estimated by subtracting the remaining glucose concentration from the starting concentration, and dividing by the time cells was under treatment condition.

### Phos-Tag electrophoresis and western blot

All samples for western blot experiments were collected from cells cultured in 6 well-plate, at 80% confluency. Samples were lysed with ice-cold RIPA buffer. For Phos-Tag™ gel electrophoresis, we used SuperSep™ Phos-tag™ Precast Gels (Wako; 195-17991). Samples were loaded at 3 ug/lane, as measured by BCA protein assay (Thermo Scientific 23225).

The electrophoresis running buffer was Tris-Glycine-SDS solution (25 mM Tris, 192 mM Glycine, 0.1% SDS, pH 8.3), supplied with 1.25 mM sodium bisulfite immediately before electrophoresis. Electrophoresis was performed at 100V, constant voltage for 3 hours at 4°C. After electrophoresis was completed, gels were washed in methanol-free transfer buffer (25 mM Tris, 192 mM Glycine, pH 8.3, 10 mM EDTA) for 3 times, 10 minutes each in order to remove bivalent cations that would immobilize phosphorylated proteins in the gel. Then gel was equilibrated in transfer buffer (25 mM Tris, 192 mM Glycine, pH 8.3, 10 mM EDTA, 20% v/v Methanol) twice, 10 minutes each. Separated proteins were transferred to PVDF with wet blot transfer method at 18V, overnight at 4°C.

Following protein transfer, membranes were stained with 3% w/v Ponceau S to validate transfer efficiency, then thoroughly destained with Milli-Q water and 0.1%PBST (10 mM Tris–HCl (pH 7.5), 100 mM NaCl, and 0.10% v/v Tween-20). Non-specific antibody binding was blocked by incubating membranes in Odyssey blocking buffer (Licor; 927-40000) for 1 hour at room temperature. Primary antibodies (Rabbit Anti-GFP, CST 2956) were diluted to 1:1000 in blocking buffer and incubated with the membrane overnight at 4°C. Following extensive washing in 0.1%PBST (3 times, 10 minutes each), membranes were incubated with diluted IRDye 800CW (Licor; 926-32211) secondary antibodies for 1 hour, at room temperature. After washing in 0.1%BST (3 times, 10 minutes each) immunoreactive bands were recorded with an Odyssey CLx imaging system. Protein identity was based on immunoreactivity.

For western blot quantification, we manually segment protein band using ImageJ and used average band intensity as representative parameter. AMPKAR2 phosphorylation fraction was calculated by computing the ratio of phosphorylated band over summation of phosphorylated and unphosphorylated bands.

### Luminescence ATP determination

ATP concentration for bulk cell populations was determined using an ATP determination kit (Thermo Fisher, A22066), using protocal provided with the kit with minor modification as follows. Cells were plated in 96-well plate at 25000 cell/well 1 day before the experiment and treated as previously described for live-cell microscopy. Samples were collected at indicated time points by incubation with Trichloracetic acid (TCA), final concentration 2.5% v/v, at 4° C for 30 minutes. After cell lysis, samples were diluted five-fold to minimize TCA concentration (now 0.5% v/v). 10 µl of diluted sample was added to 90 µl reaction solution (see product manual), in 96-well plate assay plate (Corning 3603) followed by incubation for 15 minutes at room temperature. Luminescence was monitored by microplate reader (Molecular Device, SpectraMax M5) at 560 nM, room temperature.

### Replicates and Statistical test

Numbers of independent replicates are indicated in each figure legend as ‘N’; we define ‘independent replicate’ here as a complete, separate performance of a time-lapse imaging experiment with similar culture and treatment conditions, beginning from the plating of cells from bulk culture on an imaging plate and occurring on different days from other replicates. For all experiments shown, a minimum of 300 cells was imaged and tracked for each condition. Unless noted otherwise, where single-cell recordings are shown, the displayed cells were chosen by random number generation in MATLAB with a threshold for minimum tracking time to eliminate cells in which recording was terminated prematurely due to failure of the tracking algorithm. The chosen tracks were manually verified to be representative of successfully tracked cells and consistent with the overall range of cell behaviors. Cell recordings determined by manual inspection to have poor tracking or quantification accuracy were discarded.

### Sample preparation for GC-TOF analysis

For GCMS analysis, cells were plated in 10 cm plates at 10^7^ cells per plate. After incubation overnight, the growth medium was replaced with 10 ml of ‘DMEM/F12 experiment media 1’ supplied with 17 mM glucose. After 4 hours of incubation, cells were treated with oligomycin 1.8 μg/ml. Samples were quenched by immediately replacing the media with 1 ml of pre-chilled, degassed 3:3:2 v/v acetonitrile:isopropanol:water (Fisher) at 0,30,60,150 and 270 minutes following oligomycin, representing the average first peak, trough, and second peak of the AMPKAR2^PHOS^ response to ETCi. After quenching, samples were flash-frozen in liquid nitrogen and stored in −80°C freezer.

Prior to GC-TOF analysis, all samples were thawed at room temperature and centrifuged at 14,000 rcf. Supernatants were removed and samples evaporated to dryness using a CentrVap. To remove membrane lipids and triglycerides, dried samples were resuspended with 1:1 v/v acetonitrile:water, decanted and evaporated to dryness using a CentrVap. Internal standards, C8–C30 fatty acid methyl esters (FAMEs), were added to samples and derivatized with methoxyamine hydrochloride in pyridine followed by MSTFA (Sigma-Aldrich 69479) for trimethylsilylation of acidic protons. Derivatized samples were subsequently submitted for analysis by GC-TOFMS.

### Chromatographic and mass spectrometric conditions for GC-TOF analysis

Primary metabolite data was collected using a Leco Pegasus IV time of flight (TOF) MS (Leco Corporation) coupled to an Agilent 6890 GC (Agilent Technologies) equipped with a 30 m long 0.25 mm id Rtx5Sil-MS column (30 m × 0.25 mm; 0.25 µm phase) and a Gerstel MPS2 automatic liner exchange system (Gerstel GMBH & Co. KG). The chromatographic gradient used a constant flow of 1 ml/min, and an oven temperature ramping from 50°C for to 330°C over 22 minutes. Mass spectrometry data were collected using 1525 V detector voltage at m/z 85–500 with 17 spectra/sec, electron ionization at −70 eV and an ion source temperature of 250°C. QC injections, blanks and pooled human plasma were used for quality assurance throughout the run. Data were processed by BinBase (Fiehn et al., 2005) for deconvolution, peak picking, filtering, and metabolite identifications.

### GC-TOF data analysis

Peak heights of each metabolite were used for further statistical analysis. First, data were normalized by using the sum of the knowns, or mTIC normalization, to scale each sample. Peak heights were then submitted using R to DeviumWeb (v0.3.2). The data were normalized further by log transformation and Pareto scaling. ANOVA analysis was performed with FDR correction and Tukey post hoc testing with alpha of 0.05.

### Cell line Table

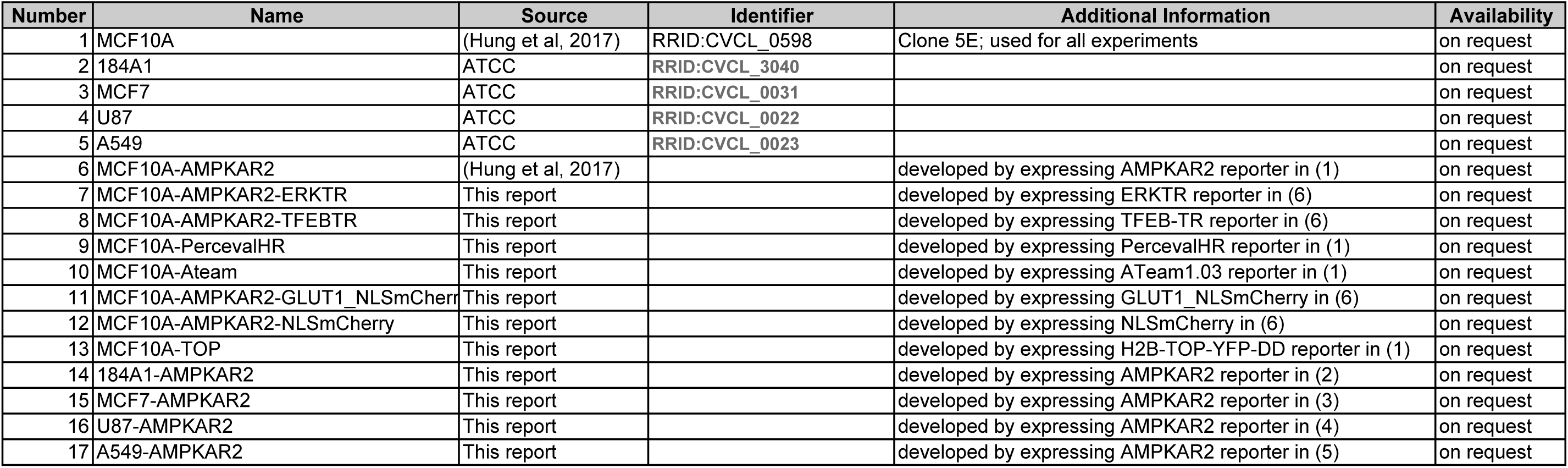

### Inhibitor List

| **Name** | **Solvent** | **Vendor** | **Catalog number** | **Remarks** |
| --- | --- | --- | --- | --- |
| Oligomycin A | DMSO | Sigma | 75351 |  |
| Antimycin A | DMSO | Sigma | A8674 |  |
| Rotenone | DMSO | Sigma | 45656 |  |
| FCCP | DMSO | Sigma | C2920 |  |
| Torin2 | DMSO | Selleckchem | S2817 |  |
| MK2206 | DMSO | Selleckchem | S1078 |  |
| Rapalink | DMSO | - | (Fan et al., 2017) | A gift from Shokat Lab |
| Torin1 | DMSO | Selleckchem | S2827 |  |
| 2DG | Water | Sigma | D8375 |  |
| 3PO | DMSO | Selleckchem | S7639 |  |
| BU99006 | DMSO | Santa Cruz Biotech | SC-300307 |  |
| Etomoxir | DMSO | Sigma | E1905 |  |
| 6AN | Water | Cayman Chemical | 10009315 |  |
| Cycloheximide | DMSO | EMD millipore | 239763 |  |
| Bafilomycin A | Water | Cayman Chemical | 11038 |  |
| SBI0206965 | DMSO | Sigma | SML1540 |  |
| PF0175157 | DMSO | Sigma | PZ0299 |  |
| Geneticin | Water | Sigma | A1720 |  |
| Torin1 | DMSO | Selleckchem | S2827 |  |

### Media table

**Media table****DMEM/F12 growth media**
| Component | Stock Concentration | Volume | Unit | Solvent | Vendor | Catalog number | Final Concentration |
| --- | --- | --- | --- | --- | --- | --- | --- |
| DMEM/F2 | - | 500 | ml | - | Gibco | 11320-033 | - |
| Horse Serum | - | 25 | ml | - | Invitrogen | 16050-122 | 5% |
| EGF | 100 ug/ml | 100 | ul | Sterile water | Peptotech | AF-100-15 | 20 ng/ml |
| Hydrocortisone | 1 mg/ml | 250 | ul | 200 proof ethanol | Sigma | H0888 | 0.5 mg/ml |
| Cholera toxin | 1 mg/ml | 50 | ul | Sterile water | Sigma | C8052 | 100 ng/ml |
| Insulin | 10 mg/ml | 500 | ul | - | Sigma | I9278 | 10 ug/ml |

**DMEM growth media**
| Component | Stock Concentration | Volume | Unit | Solvent | Vendor | Catalog number | Final Concentration |
| --- | --- | --- | --- | --- | --- | --- | --- |
| DMEM | - | 500 | ml | - | Gibco | 11965-092 | - |
| Fetal bovine serum | - | 50 | ml | - | Gemini bio products | 100-106 | 10% |

**Imaging medium 1**
| Component | Stock Concentration | Volume | Unit | Solvent | Vendor | Catalog number | Final Concentration |
| --- | --- | --- | --- | --- | --- | --- | --- |
| Imaging base-DMEM/F12 | - | 500 | ml | - | Gibco | Custom; equivalent to Gibco 11320-033 lacking glucose, glutamine, pyruvate, riboflavin, folic acid, and phenol red | - |
| BSA | 2% w/v | 25 | ml | Sterile water | Invitrogen | 16050-122 | 0.1% w/v |
| Hydrocortisone | 1 mg/ml | 250 | ul | 200 proof ethanol | Sigma | H0888 | 0.5 mg/ml |
| Cholera toxin | 1 mg/ml | 50 | ul | Sterile water | Sigma | C8052 | 100 ng/ml |
| Penicillin-Streptomycin | 10000 U/ml | 5 | ml | - | Gibco | 15140122 | 100 U/ml |
| EGF | 100 ug/ml | 100 | ul | Sterile water | Peptotech | AF-100-15 | 20 ng/ml |
| Insulin | 10 mg/ml | 500 | ul | - | Sigma | I9278 | 10 ug/ml |

**Imaging medium 2**
| Component | Stock Concentration | Volume | Unit | Solvent | Vendor | Catalog number | Final Concentration |
| --- | --- | --- | --- | --- | --- | --- | --- |
| Imaging base-DMEM/F12 | - | 500 | ml | - | Gibco | Custom; equivalent to Gibco 11320-033 lacking glucose, glutamine, pyruvate, riboflavin, folic acid, and phenol red | - |
| BSA | 2% w/v | 25 | ml | Sterile water | Invitrogen | 16050-122 | 0.1% w/v |
| Penicillin-Streptomycin | 10000 U/ml | 5 | ml | - | Gibco | 15140122 | 100 U/ml |

---

**Supplemental Figure S1.**
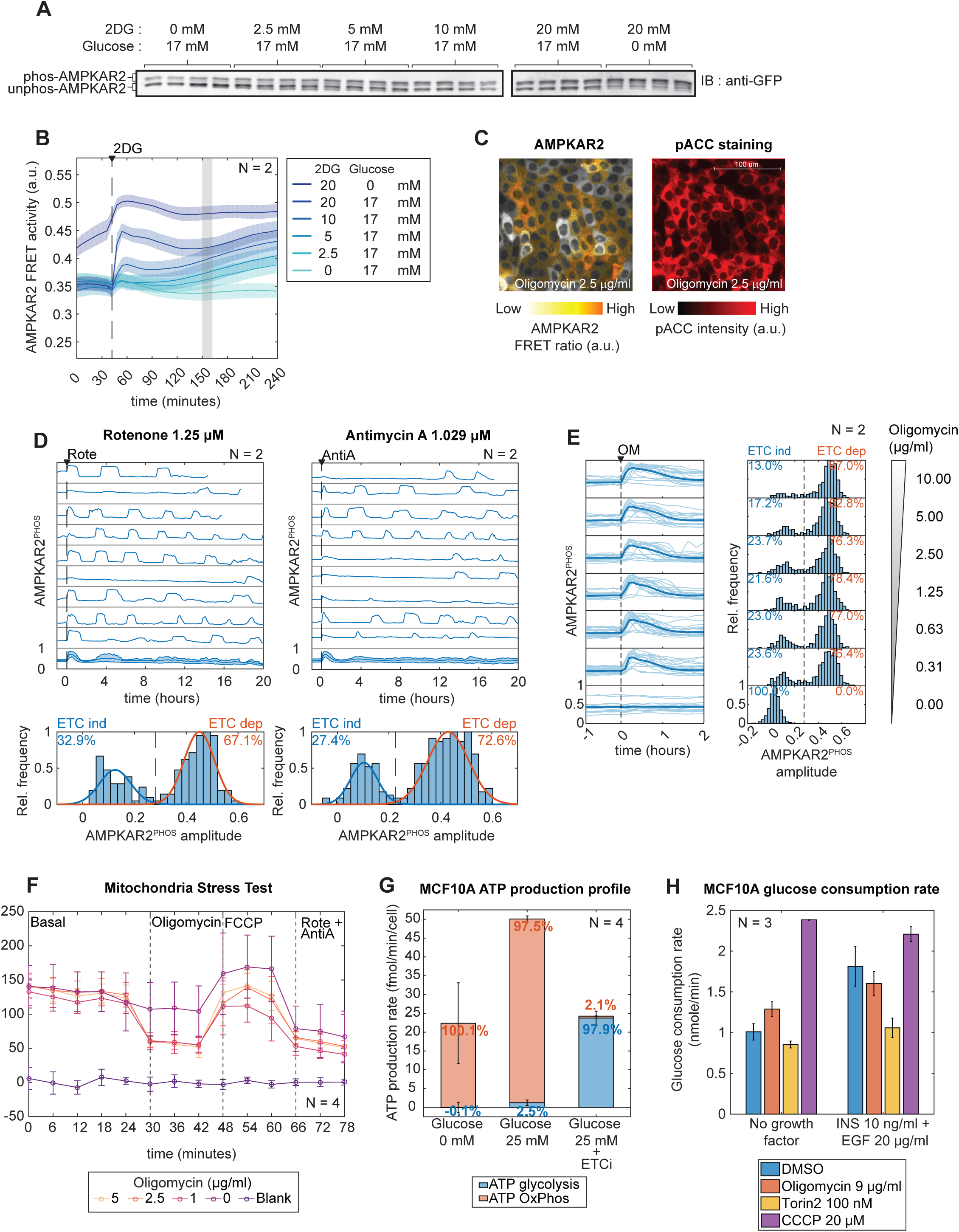
Related to Figure 1. A: Measurement of AMPKAR2 phosphorylation status. Phos-tag gel electrophoresis was used to separate phosphorylated and unphosphorylated forms of the reporter (upper and lower bands, respectively), with anti-GFP used to detect both forms. A range of different AMPK activities were induced by varying glucose and 2-deoxyglucose (2-DG). B: Population average AMPKAR2 FRET measurements the under indicated concentrations of glucose and 2-DG. The shaded area represents the time at which samples were collected for phos-tag gel electrophoresis and further immunoblot for calibration of AMPKAR2 reporter. FRET activity in shaeded area is use for scatter plot in Figure 1B. C: Representative image of AMPKAR2 FRET activity (shown as a ratiometric image) and the corresponding pACC immunofluorescence pattern after the same cells were fixed and stained. D: Upper panels: sample of AMPKAR2 measurement after treated with Rotenone 1.25 µM (left panel) or Antimycin A 1.029 µM (right panel). Each subplot represents a single cell measurement, with population average and interquartile range in the bottom subplot. Lower panels: Histogram of AMPKAR2 response amplitude after treatment with Rotenone 1.25 µM (left panel) or Antimycin A 1.029 µM (right panel). Blue and orange lines are fitted Gaussian distributions. The dashed line is defined by the intersection between distributions and used as cutoff for ETC-ind and ETC-dep cells. E: Left panel: sample single-cell activities of each reporter after cells were cultured in media containing 17 mM glucose and then treated with oligomycin (OM) at the concentrations indicated. Dark lines – AMPKAR2^PHOS^ population average. Light lines – single cell AMPKAR2^PHOS^ measurement. Right panel: Histograms of response amplitude of AMPKAR2 with treatment conditions corresponding to left panel. F: Oxygen consumption rate of MCF10A-AMPKAR2 cell line measured by Seahorse XF assay. Cells were treated with oligomycin as indicated followed by FCCP 5 µM and combination of rotenone 2.5 µM and antimycin A 2.5 µM to completely inhibit cellular respiration. ‘Blank’ indicates measurement from well without any cells G: Estimated ATP production by glycolysis and OxPhos in MCF10A cell line under indicated condition. Error bars represent the standard deviations of four replicates. H: Estimated glucose uptake rate of MCF10A cells at condition indicated. Error bars represent the standard deviation of three replicates.

**Supplemental Figure S2.**
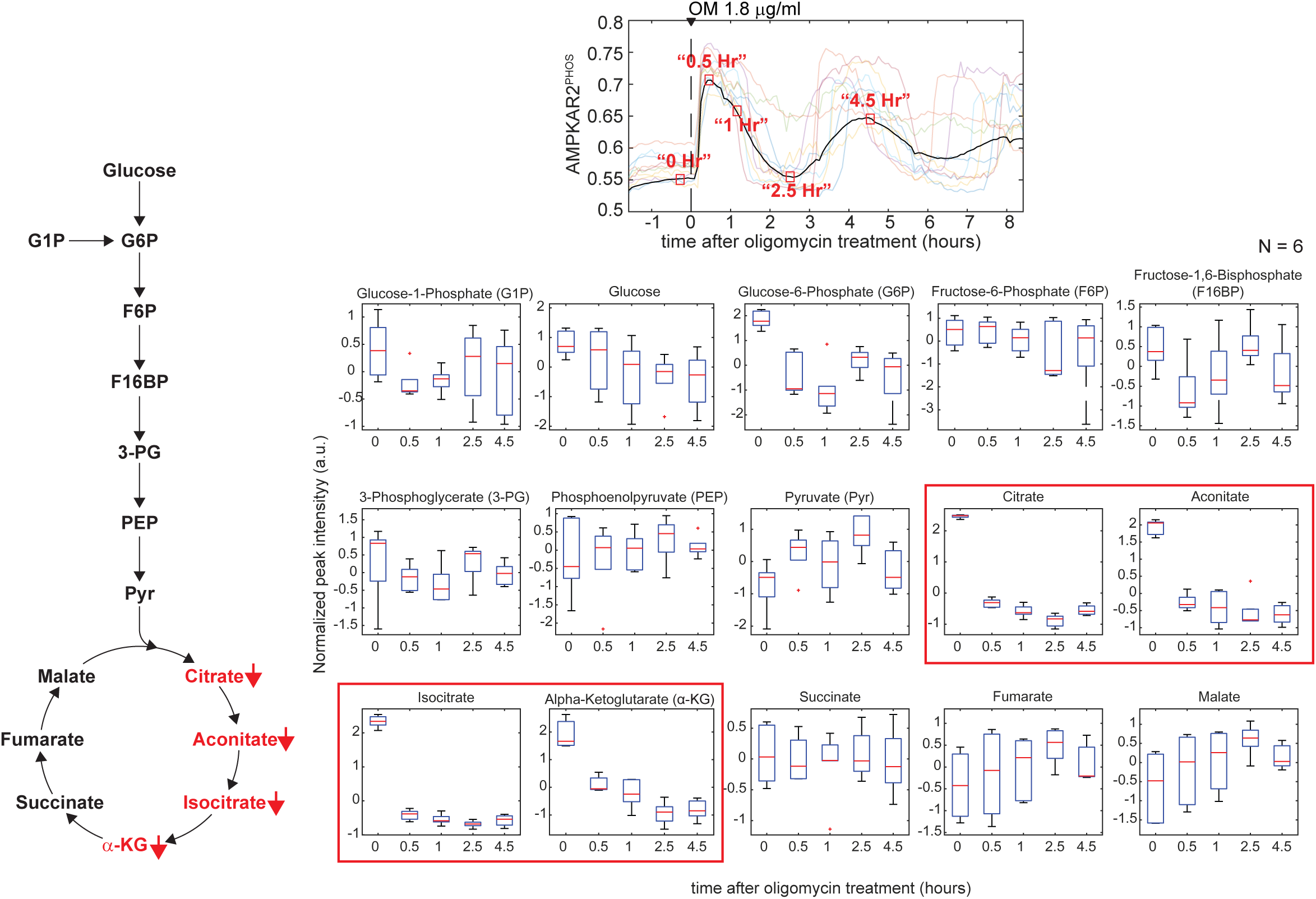
Related to Figure 1. Upper panel: schematic representation of population average AMPKAR2^PHOS^ after oligomycin treatment. Red boxes indicate time points at which samples were collected for mass spectrometry analysis. Left panel: schematic of metabolites in glycolysis and TCA cycle that were identified from mass spectrometry analysis. Right panel: box plots of relative concentration of all metabolites identified by mass spectrometry. Red font and boxes show metabolites that were significantly reduced after ETCi treatment.

**Supplemental Figure S3.**
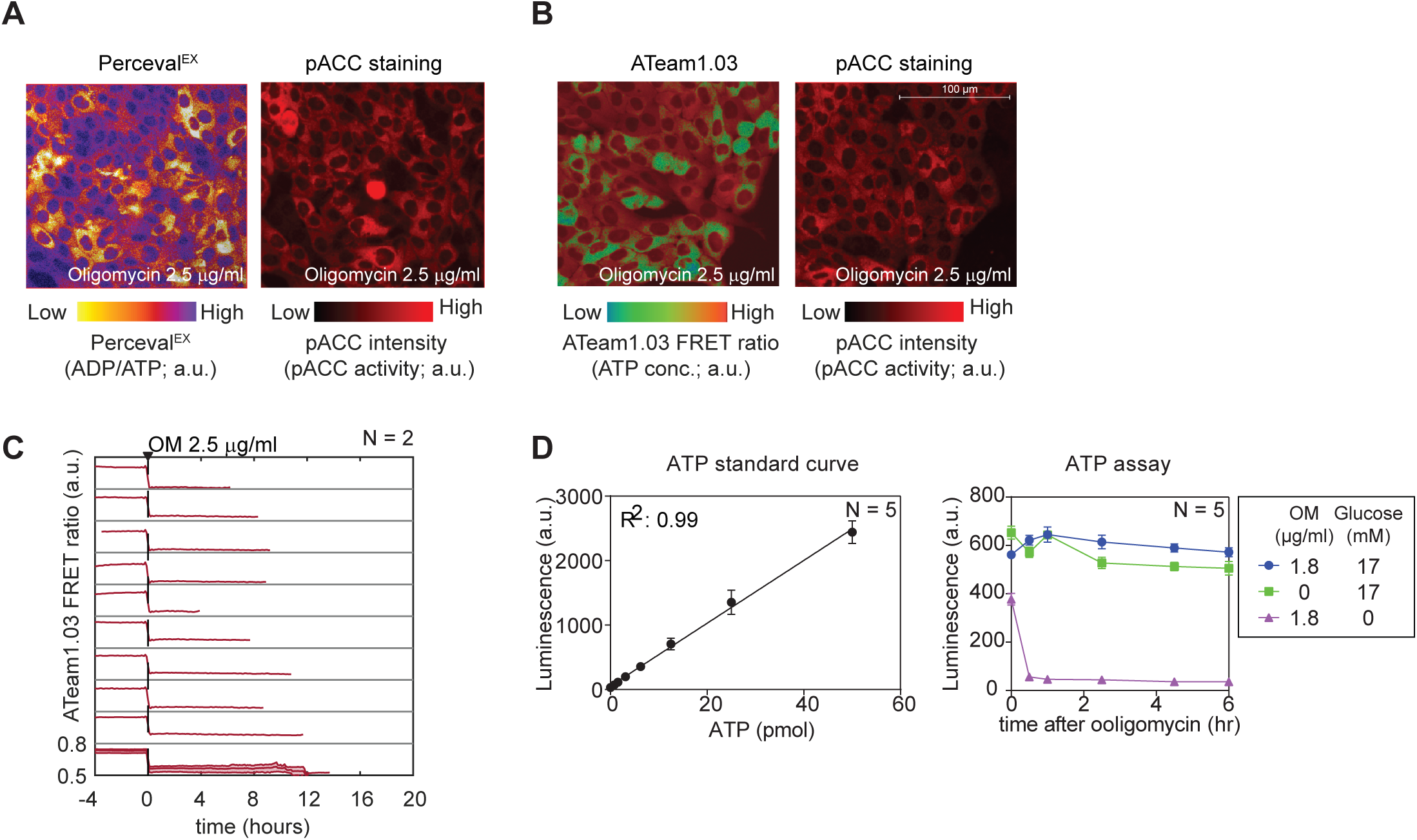
Related to Figure 2. A and B: Sample image of Perceval^EX^ and ATeam1.03 FRET activity and their corresponding pACC staining pattern in response to 2.5 µg/ml oligomycin. Reporter images were acquired in live cells at 30 minutes after oligomycin treatment. Cells were then immediately fixed and stained for pACC. C: ATeam1.03 measurements after cells were treated with 2.5 µg/ml oligomycin in the absence of glucose. Each subplot represents a single cell measurement, with population average and interquartile range in the bottom subplot. D: Left panel: standard curve of bulk ATP measurement. Right panel: Bulk ATP measurement of MCF10A cell line under condition indicated.

**Supplemental Figure S4.**
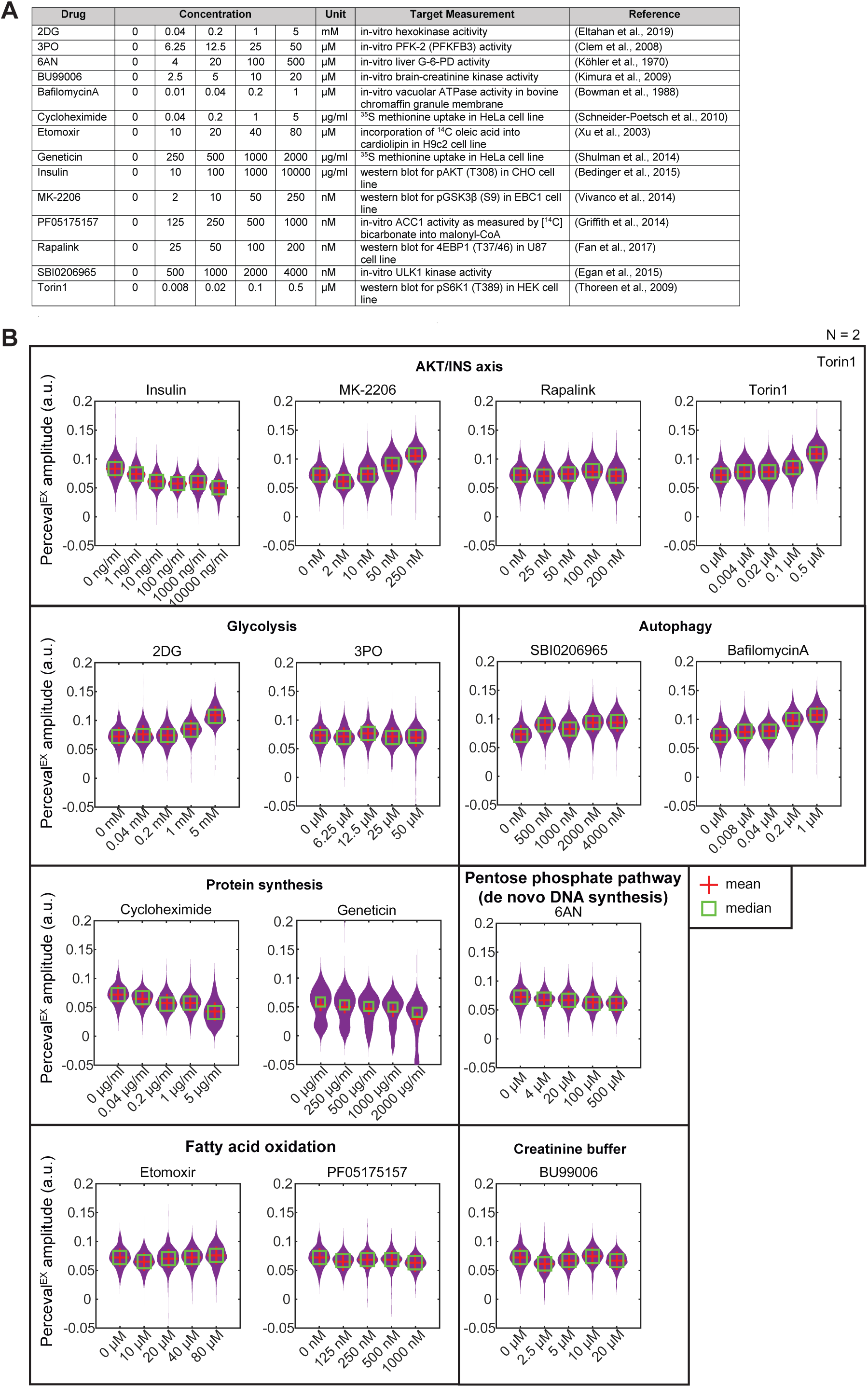
Related to Figure 4. A: Table showing inhibitors used in Figure 4E-4G, and their corresponding concentration with published target activity measurements. B: Violin plots of the distributions of Perceval^EX^ amplitude when MCF10A cells are pretreated with inhibitors as indicated for 30 minutes, followed by treatment with oligomycin.

**Supplemental Figure S5.**
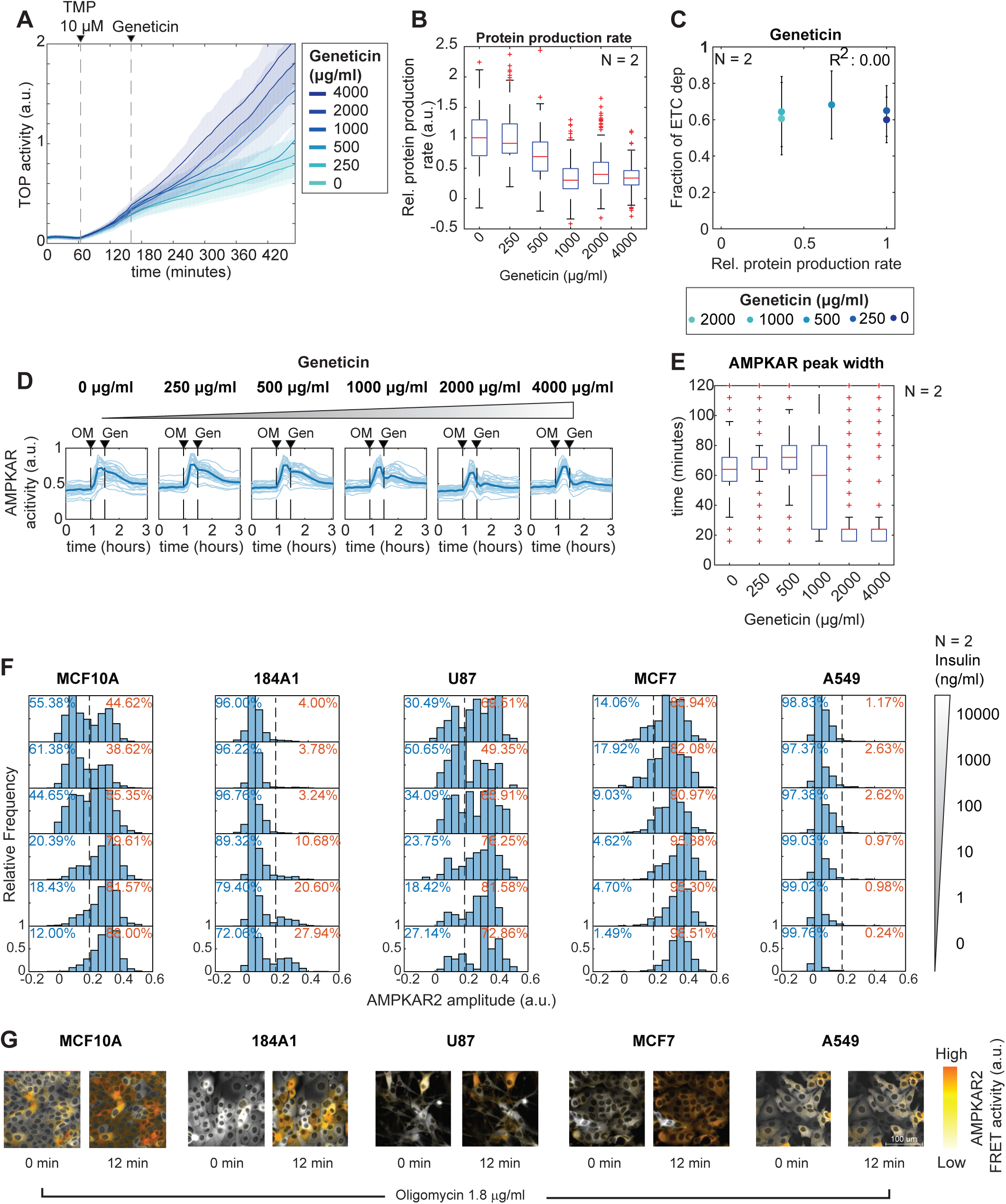
Related to Figure 5. A: Representative mean translation reporter results for a concentration series of geneticin. Shaded areas show interquartile ranges. B: Boxplot of calculated single-cell relative protein production rates for the indicated dose of geneticin. Each box represents the distribution of 400 cells. C: Scatter plot showing popultion average relative protein production rate as measured by H2B-TOP-YFP-DD when treated with geneticin and their corresponding fraction of cells that become ETC dep after challenged with oligomcyin. D: Sample of AMPKAR2^PHOS^ when treated with oligomycin and then with geneticin at the doses indicated. OM – oligomycin; Gen – genetcin; Dark lines – AMPKAR2^PHOS^ population average; Light lines – single cell AMPKAR2^PHOS^ measurement E: Boxplot of single cell AMPKAR2^PHOS^ pulse widths after geneticin treatment. Pulse widths were calculated as the time at which AMPKAR^PHOS^ decreased to 50% of the maximum value for each cell following treatment with geneticin. F: Histogram of AMPKAR2^PHOS^ amplitude reponse to oligomycin 1.8 µg/ml treatament in MCF10A, 184A, U87, MCF7 and A549 cell line under insulin concentration as indicated. G: Sample image of AMPKAR2 response when treated by oligomycin.

**Supplemental Figure S6.**
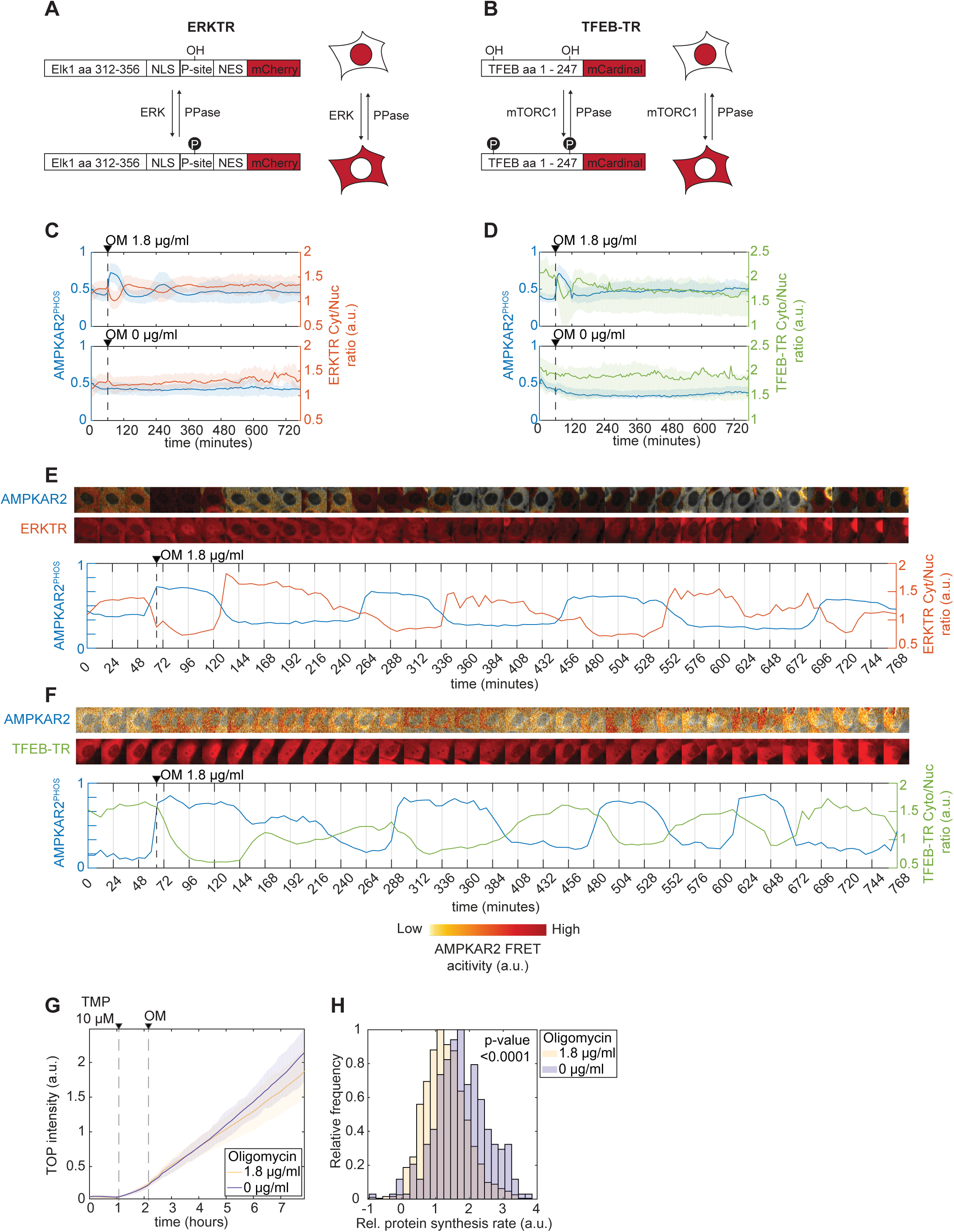
Related to Figure 6. A and B: Schematic of ERKTR and TFEB-TR reporters. C: Population average of AMPKAR2^PHOS^ (blue) and ERKTR (orange) activity after oligomcyin (upper panel) or vehicle (lower panel) treatment. D: Population average of AMPKAR2^PHOS^ (blue) and TFEB-TR (green) activity after oligomcyin (upper panel) or vehicle (lower panel) treatment. E: Sample images and single cell tracing of AMPKAR2^PHOS^ and ERKTR (orange) activity after oligomcyin. E: Sample images and single cell tracing of AMPKAR2^PHOS^ and TFEB-TR (green) activity after oligomcyin.

## References

Ahn, E., Kumar, P., Mukha, D., Tzur, A., and Shlomi, T. (2017). Temporal fluxomics reveals oscillations in TCA cycle flux throughout the mammalian cell cycle. Mol Syst Biol 13, 953.

Argüello-Miranda, O., Liu, Y., Wood, N.E., Kositangool, P., and Doncic, A. (2018). Integration of Multiple Metabolic Signals Determines Cell Fate Prior to Commitment. Mol Cell.

Bass, J., and Takahashi, J.S. (2010). Circadian integration of metabolism and energetics. Science 330, 1349–1354.

Bedinger, D.H., Goldfine, I.D., Corbin, J.A., Roell, M.K., and Adams, S.H. (2015). Differential pathway coupling of the activated insulin receptor drives signaling selectivity by XMetA, an allosteric partial agonist antibody. J Pharmacol Exp Ther 353, 35–43.

Bowman, E.J., Siebers, A., and Altendorf, K. (1988). Bafilomycins: a class of inhibitors of membrane ATPases from microorganisms, animal cells, and plant cells. Proc Natl Acad Sci U S A 85, 7972–7976.

Buttgereit, F., and Brand, M.D. (1995). A hierarchy of ATP-consuming processes in mammalian cells. Biochem J 312 (*Pt 1*), 163–167.

Cai, L., and Tu, B.P. (2012). Driving the cell cycle through metabolism. Annu Rev Cell Dev Biol 28, 59–87.

Cavanaugh, J.E. (1997). Unifying the derivations for the Akaike and corrected Akaike information criteria. Statistics & Probability Letters 33, 201–208.

Clem, B., Telang, S., Clem, A., Yalcin, A., Meier, J., Simmons, A., Rasku, M.A., Arumugam, S., Dean, W.L., Eaton, J., et al. (2008). Small-molecule inhibition of 6-phosphofructo-2-kinase activity suppresses glycolytic flux and tumor growth. Mol Cancer Ther 7, 110–120.

Cokorinos, E.C., Delmore, J., Reyes, A.R., Albuquerque, B., Kjobsted, R., Jorgensen, N.O., Tran, J.L., Jatkar, A., Cialdea, K., Esquejo, R.M., et al. (2017). Activation of Skeletal Muscle AMPK Promotes Glucose Disposal and Glucose Lowering in Non-human Primates and Mice. Cell Metab 25, 1147–1159 e1110.

DeBerardinis, R.J., Lum, J.J., Hatzivassiliou, G., and Thompson, C.B. (2008). The biology of cancer: metabolic reprogramming fuels cell growth and proliferation. Cell Metab 7, 11–20.

Debnath, J., Muthuswamy, S.K., and Brugge, J.S. (2003). Morphogenesis and oncogenesis of MCF-10A mammary epithelial acini grown in three-dimensional basement membrane cultures. Methods 30, 256–268.

Egan, D.F., Chun, M.G.H., Vamos, M., Zou, H., Rong, J., Miller, C.J., Lou, H.J., Raveendra-Panickar, D., Yang, C.-C., Sheffler, D.J., et al. (2015). Small Molecule Inhibition of the Autophagy Kinase ULK1 and Identification of ULK1 Substrates. Mol Cell 59, 285–297.

Eltahan, R., Guo, F., Zhang, H., and Zhu, G. (2019). The Action of the Hexokinase Inhibitor 2-deoxy-d-glucose on Cryptosporidium parvum and the Discovery of Activities against the Parasite Hexokinase from Marketed Drugs. J Eukaryot Microbiol 66, 460–468.

Fan, J., Kamphorst, J.J., Mathew, R., Chung, M.K., White, E., Shlomi, T., and Rabinowitz, J.D. (2013). Glutamine-driven oxidative phosphorylation is a major ATP source in transformed mammalian cells in both normoxia and hypoxia. Mol Syst Biol 9, 712.

Fan, Q., Aksoy, O., Wong, R.A., Ilkhanizadeh, S., Novotny, C.J., Gustafson, W.C., Truong, A.Y.-Q., Cayanan, G., Simonds, E.F., Haas-Kogan, D., et al. (2017). A Kinase Inhibitor Targeted to mTORC1 Drives Regression in Glioblastoma. Cancer cell 31, 424–435.

Fendt, S.-M., Bell, E.L., Keibler, M.A., Olenchock, B.A., Mayers, J.R., Wasylenko, T.M., Vokes, N.I., Guarente, L., Vander Heiden, M.G., and Stephanopoulos, G. (2013). Reductive glutamine metabolism is a function of the α-ketoglutarate to citrate ratio in cells. Nat Commun 4, 2236.

Fiehn, O., Wohlgemuth, G., and Sholz, M. (2005). Setup and annotation of metabolomic experiments by integrating biological and mass spectrometric metadata. Dils LNBI 3615, 224–239.

Gatenby, R.A., and Gillies, R.J. (2004). Why do cancers have high aerobic glycolysis? Nat Rev Cancer 4, 891–899.

Gillies, T.E., Pargett, M., Minguet, M., Davies, A.E., and Albeck, J.G. (2017). Linear Integration of ERK Activity Predominates over Persistence Detection in Fra-1 Regulation. Cell Systems 5, 549–563.e545.

Gowans, G.J., Hawley, S.A., Ross, F.A., and Hardie, D.G. (2013). AMP is a true physiological regulator of AMP-activated protein kinase by both allosteric activation and enhancing net phosphorylation. Cell Metab 18, 556–566.

Grassian, A.R., Coloff, J.L., and Brugge, J.S. (2011). Extracellular matrix regulation of metabolism and implications for tumorigenesis. Cold Spring Harb Symp Quant Biol 76, 313–324.

Griffith, D.A., Kung, D.W., Esler, W.P., Amor, P.A., Bagley, S.W., Beysen, C., Carvajal-Gonzalez, S., Doran, S.D., Limberakis, C., Mathiowetz, A.M., et al. (2014). Decreasing the rate of metabolic ketone reduction in the discovery of a clinical acetyl-CoA carboxylase inhibitor for the treatment of diabetes. J Med Chem 57, 10512–10526.

Gwinn, D.M., Shackelford, D.B., Egan, D.F., Mihaylova, M.M., Mery, A., Vasquez, D.S., Turk, B.E., and Shaw, R.J. (2008). AMPK phosphorylation of raptor mediates a metabolic checkpoint. Mol Cell 30, 214–226.

Hackett, S.R., Zanotelli, V.R.T., Xu, W., Goya, J., Park, J.O., Perlman, D.H., Gibney, P.A., Botstein, D., Storey, J.D., and Rabinowitz, J.D. (2016). Systems-level analysis of mechanisms regulating yeast metabolic flux. Science 354.

Han, K., Jaimovich, A., Dey, G., Ruggero, D., Meyuhas, O., Sonenberg, N., and Meyer, T. (2014a). Parallel measurement of dynamic changes in translation rates in single cells. Nat Methods 11, 86–93.

Han, K., Jaimovich, A., Dey, G., Ruggero, D., Meyuhas, O., Sonenberg, N., and Meyer, T. (2014b). Parallel measurement of dynamic changes in translation rates in single cells. Nat Methods 11, 86–93.

Hardie, D.G. (2014). AMPK—Sensing Energy while Talking to Other Signaling Pathways. Cell Metab 20, 939–952.

Hardie, D.G., Ross, F.A., and Hawley, S.A. (2012). AMPK: a nutrient and energy sensor that maintains energy homeostasis. Nat Rev Mol Cell Biol 13, 251–262.

Hoppe, A., Christensen, K., and Swanson, J.A. (2002). Fluorescence Resonance Energy Transfer-Based Stoichiometry in Living Cells. Biophysical Journal 83, 3652–3664.

Hung, Y.P., Teragawa, C., Kosaisawe, N., Gillies, T.E., Pargett, M., Minguet, M., Distor, K., Rocha-Gregg, B.L., Coloff, J.L., Keibler, M.A., et al. (2017a). Akt regulation of glycolysis mediates bioenergetic stability in epithelial cells. Elife 6.

Hung, Y.P., Teragawa, C., Kosaisawe, N., Gillies, T.E., Pargett, M., Minguet, M., Distor, K., Rocha-Gregg, B.L., Coloff, J.L., Keibler, M.A., et al. (2017b). Akt regulation of glycolysis mediates bioenergetic stability in epithelial cells. eLife 6, 1–25.

Imamura, H., Huynh Nhat, K.P., Togawa, H., Saito, K., Iino, R., Kato-Yamada, Y., Nagai, T., and Noji, H. (2009a). Visualization of ATP levels inside single living cells with fluorescence resonance energy transfer-based genetically encoded indicators. Proceedings of the National Academy of Sciences 106, 15651–15656.

Imamura, H., Nhat, K.P.H., Togawa, H., Saito, K., Iino, R., Kato-Yamada, Y., Nagai, T., and Noji, H. (2009b). Visualization of ATP levels inside single living cells with fluorescence resonance energy transfer-based genetically encoded indicators. Proc Natl Acad Sci U S A 106, 15651–15656.

Inoki, K., Zhu, T., and Guan, K.L. (2003). TSC2 mediates cellular energy response to control cell growth and survival. Cell 115, 577–590.

Janes, K.A., Wang, C.-C., Holmberg, K.J., Cabral, K., and Brugge, J.S. (2010). Identifying single-cell molecular programs by stochastic profiling. Nat Methods 7, 311–317.

Jaqaman, K., Loerke, D., Mettlen, M., Kuwata, H., Grinstein, S., Schmid, S.L., and Danuser, G. (2008). Robust single-particle tracking in live-cell time-lapse sequences. Nature Methods 5, 695–702.

Jeon, S.-M., Chandel, N.S., and Hay, N. (2012). AMPK regulates NADPH homeostasis to promote tumour cell survival during energy stress. Nature 485, 661–665.

Kimura, A., Tyacke, R.J., Robinson, J.J., Husbands, S.M., Minchin, M.C.W., Nutt, D.J., and Hudson, A.L. (2009). Identification of an imidazoline binding protein: creatine kinase and an imidazoline-2 binding site. Brain Res 1279, 21–28.

Köhler, E., Barrach, H.-J., and Neubert, D. (1970). Inhibition of NADP dependent oxidoreductases by the 6-aminonicotinamide analogue of NADP. FEBS Lett 6, 225–228.

Konagaya, Y., Terai, K., Hirao, Y., Takakura, K., Imajo, M., Kamioka, Y., Sasaoka, N., Kakizuka, A., Sumiyama, K., Asano, T., et al. (2017). A Highly Sensitive FRET Biosensor for AMPK Exhibits Heterogeneous AMPK Responses among Cells and Organs. Cell Rep 21, 2628–2638.

Li, L., Friedrichsen, H.J., Andrews, S., Picaud, S., Volpon, L., Ngeow, K., Berridge, G., Fischer, R., Borden, K.L.B., Filippakopoulos, P., et al. (2018). A TFEB nuclear export signal integrates amino acid supply and glucose availability. Nat Commun 9, 2685.

Lin, S.-C., and Hardie, D.G. (2017). AMPK: Sensing Glucose as well as Cellular Energy Status. Cell Metab.

Lunt, S.Y., and Vander Heiden, M.G. (2011). Aerobic glycolysis: meeting the metabolic requirements of cell proliferation. Annu Rev Cell Dev Biol 27, 441–464.

Mookerjee, S.A., Gerencser, A.A., Nicholls, D.G., and Brand, M.D. (2017a). Quantifying intracellular rates of glycolytic and oxidative ATP production and consumption using extracellular flux measurements. J Biol Chem 292, 7189–7207.

Mookerjee, S.A., Gerencser, A.A., Nicholls, D.G., and Brand, M.D. (2017b). Quantifying intracellular rates of glycolytic and oxidative ATP production and consumption using extracellular flux measurements. J Biol Chem 292, 7189–7207.

Myers, R.W., Guan, H.P., Ehrhart, J., Petrov, A., Prahalada, S., Tozzo, E., Yang, X., Kurtz, M.M., Trujillo, M., Gonzalez Trotter, D., et al. (2017). Systemic pan-AMPK activator MK-8722 improves glucose homeostasis but induces cardiac hypertrophy. Science 357, 507–511.

O’Neill, H.M., Maarbjerg, S.J., Crane, J.D., Jeppesen, J., Jørgensen, S.B., Schertzer, J.D., Shyroka, O., Kiens, B., van Denderen, B.J., Tarnopolsky, M.A., et al. (2011). AMP-activated protein kinase (AMPK) beta1beta2 muscle null mice reveal an essential role for AMPK in maintaining mitochondrial content and glucose uptake during exercise. Proc Natl Acad Sci U S A 108, 16092–16097.

Papagiannakis, A., Niebel, B., Wit, E.C., and Heinemann, M. (2017). Autonomous Metabolic Oscillations Robustly Gate the Early and Late Cell Cycle. Mol Cell 65, 285–295.

Pargett, M., Gillies, T.E., Teragawa, C.K., Sparta, B., and Albeck, J.G. (2017). Single-Cell Imaging of ERK Signaling Using Fluorescent Biosensors (Humana Press, New York, NY), pp. 35–59.

Regot, S., Hughey, J.J., Bajar, B.T., Carrasco, S., and Covert, M.W. (2014). High-sensitivity measurements of multiple kinase activities in live single cells. Cell 157, 1724–1734.

Rodrik-Outmezguine, V.S., Okaniwa, M., Yao, Z., Novotny, C.J., McWhirter, C., Banaji, A., Won, H., Wong, W., Berger, M., de Stanchina, E., et al. (2016). Overcoming mTOR resistance mutations with a new-generation mTOR inhibitor. Nature 534, 272–276.

Roy, S., Leidal, A.M., Ye, J., Ronen, S.M., and Debnath, J. (2017). Autophagy-Dependent Shuttling of TBC1D5 Controls Plasma Membrane Translocation of GLUT1 and Glucose Uptake. Mol Cell 67, 84–95.e85.

Schafer, Z.T., Grassian, A.R., Song, L., Jiang, Z., Gerhart-Hines, Z., Irie, H.Y., Gao, S., Puigserver, P., and Brugge, J.S. (2009). Antioxidant and oncogene rescue of metabolic defects caused by loss of matrix attachment. Nature 461, 109–113.

Schneider-Poetsch, T., Ju, J., Eyler, D.E., Dang, Y., Bhat, S., Merrick, W.C., Green, R., Shen, B., and Liu, J.O. (2010). Inhibition of eukaryotic translation elongation by cycloheximide and lactimidomycin. Nat Chem Biol 6, 209–217.

Settembre, C., Zoncu, R., Medina, D.L., Vetrini, F., Erdin, S., Erdin, S., Huynh, T., Ferron, M., Karsenty, G., Vellard, M.C., et al. (2012). A lysosome-to-nucleus signalling mechanism senses and regulates the lysosome via mTOR and TFEB. EMBO J 31, 1095–1108.

Shen, C.H., Yuan, P., Perez-Lorenzo, R., Zhang, Y., Lee, S.X., Ou, Y., Asara, J.M., Cantley, L.C., and Zheng, B. (2013). Phosphorylation of BRAF by AMPK impairs BRAF-KSR1 association and cell proliferation. Mol Cell 52, 161–172.

Shi, L., and Tu, B.P. (2013). Acetyl-CoA induces transcription of the key G1 cyclin CLN3 to promote entry into the cell division cycle in Saccharomyces cerevisiae. Proc Natl Acad Sci U S A 110, 7318–7323.

Shulman, E., Belakhov, V., Wei, G., Kendall, A., Meyron-Holtz, E.G., Ben-Shachar, D., Schacht, J., and Baasov, T. (2014). Designer aminoglycosides that selectively inhibit cytoplasmic rather than mitochondrial ribosomes show decreased ototoxicity: a strategy for the treatment of genetic diseases. J Biol Chem 289, 2318–2330.

Sigal, A., Milo, R., Cohen, A., Geva-Zatorsky, N., Klein, Y., Alaluf, I., Swerdlin, N., Perzov, N., Danon, T., Liron, Y., et al. (2006a). Dynamic proteomics in individual human cells uncovers widespread cell-cycle dependence of nuclear proteins. Nat Methods 3, 525–531.

Sigal, A., Milo, R., Cohen, A., Geva-Zatorsky, N., Klein, Y., Liron, Y., Rosenfeld, N., Danon, T., Perzov, N., and Alon, U. (2006b). Variability and memory of protein levels in human cells. Nature 444, 643–646.

Silverman, S.J., Petti, A.A., Slavov, N., Parsons, L., Briehof, R., Thiberge, S.Y., Zenklusen, D., Gandhi, S.J., Larson, D.R., Singer, R.H., et al. (2010). Metabolic cycling in single yeast cells from unsynchronized steady-state populations limited on glucose or phosphate. Proc Natl Acad Sci U S A 107, 6946–6951.

Simões, R.V., Serganova, I.S., Kruchevsky, N., Leftin, A., Shestov, A.A., Thaler, H.T., Sukenick, G., Locasale, J.W., Blasberg, R.G., Koutcher, J.A., et al. (2015). Metabolic plasticity of metastatic breast cancer cells: adaptation to changes in the microenvironment. Neoplasia 17, 671–684.

Slavov, N., Macinskas, J., Caudy, A., and Botstein, D. (2011). Metabolic cycling without cell division cycling in respiring yeast. Proc Natl Acad Sci U S A 108, 19090–19095.

Sparta, B., Pargett, M., Minguet, M., Distor, K., Bell, G., and Albeck, J.G. (2015). Receptor Level Mechanisms Are Required for Epidermal Growth Factor (EGF)-stimulated Extracellular Signal-regulated Kinase (ERK) Activity Pulses. The Journal of biological chemistry 290, 24784–24792.

Spencer, S.L., Gaudet, S., Albeck, J.G., Burke, J.M., and Sorger, P.K. (2009). Non-genetic origins of cell-to-cell variability in TRAIL-induced apoptosis. Nature 459, 428–432.

Strasen, J., Sarma, U., Jentsch, M., Bohn, S., Sheng, C., Horbelt, D., Knaus, P., Legewie, S., and Loewer, A. (2018). Cell-specific responses to the cytokine TGFβ are determined by variability in protein levels. Mol Syst Biol 14, e7733.

Suzuki, T., Bridges, D., Nakada, D., Skiniotis, G., Morrison, S.J., Lin, J.D., Saltiel, A.R., and Inoki, K. (2013). Inhibition of AMPK catabolic action by GSK3. Mol Cell 50, 407–419.

Szymańska, P., Martin, K.R., MacKeigan, J.P., Hlavacek, W.S., and Lipniacki, T. (2015). Computational analysis of an autophagy/translation switch based on mutual inhibition of MTORC1 and ULK1. PLoS One 10, e0116550.

Takanaga, H., Chaudhuri, B., and Frommer, W.B. (2008). GLUT1 and GLUT9 as major contributors to glucose influx in HepG2 cells identified by a high sensitivity intramolecular FRET glucose sensor. Biochimica et Biophysica Acta (BBA) - Biomembranes 1778, 1091–1099.

Tantama, M., Martinez-Francois, J.R., Mongeon, R., and Yellen, G. (2013a). Imaging energy status in live cells with a fluorescent biosensor of the intracellular ATP-to-ADP ratio. Nat Commun 4, 2550.

Tantama, M., Ramón, M.i.-F.J., and Mongeon, R., and Yellen, G. (2013b). Imaging energy status in live cells with a fluorescent biosensor of the intracellular ATP-to-ADP ratio. Nat Commun 4, 2550.

Thoreen, C.C., Kang, S.A., Chang, J.W., Liu, Q., Zhang, J., Gao, Y., Reichling, L.J., Sim, T., Sabatini, D.M., and Gray, N.S. (2009). An ATP-competitive mammalian target of rapamycin inhibitor reveals rapamycin-resistant functions of mTORC1. J Biol Chem 284, 8023–8032.

Tsou, P., Zheng, B., Hsu, C.-H., Sasaki, A.T., and Cantley, L.C. (2011). A fluorescent reporter of AMPK activity and cellular energy stress. Cell Metab 13, 476–486.

Tu, B.P., Mohler, R.E., Liu, J.C., Dombek, K.M., Young, E.T., Synovec, R.E., and McKnight, S.L. (2007). Cyclic changes in metabolic state during the life of a yeast cell. Proc Natl Acad Sci U S A 104, 16886–16891.

Vander Heiden, M.G., and DeBerardinis, R.J. (2017). Understanding the Intersections between Metabolism and Cancer Biology. Cell 168, 657–669.

Vincent, L., and Soille, P. (1991). Watersheds in digital spaces: an efficient algorithm based on immersion simulations. IEEE Transactions on Pattern Analysis and Machine Intelligence 13, 583–598.

Viollet, B., and Foretz, M. (2016). Animal Models to Study AMPK. Exp Suppl 107, 441–469.

Vivanco, I., Chen, Z.C., Tanos, B., Oldrini, B., Hsieh, W.-Y., Yannuzzi, N., Campos, C., and Mellinghoff, I.K. (2014). A kinase-independent function of AKT promotes cancer cell survival. Elife 3.

Witczak, C.A., Sharoff, C.G., and Goodyear, L.J. (2008). AMP-activated protein kinase in skeletal muscle: from structure and localization to its role as a master regulator of cellular metabolism. Cell Mol Life Sci 65, 3737–3755.

Wollman, R. (2018). Robustness, Accuracy, and Cell State Heterogeneity in Biological Systems. Curr Opin Syst Biol 8, 46–50.

Xiao, B., Sanders, M.J., Underwood, E., Heath, R., Mayer, F.V., Carmena, D., Jing, C., Walker, P.A., Eccleston, J.F., Haire, L.F., et al. (2011). Structure of mammalian AMPK and its regulation by ADP. Nature 472, 230–233.

Xu, F.Y., Taylor, W.A., Hurd, J.A., and Hatch, G.M. (2003). Etomoxir mediates differential metabolic channeling of fatty acid and glycerol precursors into cardiolipin in H9c2 cells. J Lipid Res 44, 415–423.

Yusa, K., Zhou, L., Li, M.A., Bradley, A., and Craig, N.L. (2011). A hyperactive piggyBac transposase for mammalian applications. Proceedings of the National Academy of Sciences 108, 1531–1536.

Zhang, C.-S., Hawley, S.A., Zong, Y., Li, M., Wang, Z., Gray, A., Ma, T., Cui, J., Feng, J.-W., Zhu, M., et al. (2017). Fructose-1,6-bisphosphate and aldolase mediate glucose sensing by AMPK. Nature 548, 112–116.

